# DNA methylation Dependent Restriction of Tyrosine Hydroxylase Contributes to Pancreatic *β*-cell Heterogeneity

**DOI:** 10.1101/2022.05.06.490953

**Authors:** Nazia Parveen, Jean Kimi Wang, Supriyo Bhattacharya, Janielle Cuala, Mohan Singh Rajkumar, Xiwei Wu, Hung-Ping Shih, Senta K. Georgia, Sangeeta Dhawan

**Author notes:** These authors contributed equally (Nazia Parveen, Jean Kimi Wang).

## Abstract

The molecular and functional heterogeneity of pancreatic *β*-cells is well recognized. Pancreatic islets harbor a small subset of *β*-cells that co-express Tyrosine Hydroxylase (TH), an enzyme involved in synthesis of catecholamines that repress insulin secretion. Restriction of this sub-population within islets is essential for appropriate insulin secretion. However, the distinguishing characteristics of this subpopulation and the mechanisms that restrict TH expression in *β*-cells are not known. Here, we define the specific molecular and metabolic characteristics of the TH+ *β*-cells and show that TH expression in *β*-cells is restricted by DNA methylation patterning during *β*-cell lineage specification. Ablation of *de novo* DNA methyltransferase Dnmt3a in the pancreatic- and endocrine-progenitor lineages results in a dramatic increase in the proportion of TH+ *β*-cells, while *β*-cell specific ablation of Dnmt3a has no effect on this sub-population. We demonstrate that maintenance of *Th* promoter DNA methylation patterns is essential for its continued restriction in postnatal *β*-cells, and that loss of DNA methylation dysregulates TH expression in *β*-cells in response to chronic overnutrition, contributing to impairment of *β*-cell identity. These data highlight the essential requirement of DNA methylation patterning in regulating endocrine cell fates, and reveal a novel role of DNA methylation in *β*-cell heterogeneity.

## Introduction

Pancreatic islets comprise of several cell types including the insulin producing *β*-cells, which collectively play a central role in the regulation of glucose homeostasis. *β*-cells display molecular and functional heterogeneity, with unique *β*-cell subpopulations characterizing specific developmental and disease contexts (Pipeleers 1992; Dorrell et al. 2016; Johnston et al. 2016; Rodnoi et al. 2017; Benninger and Kravets 2021). Such heterogeneity can subserve robust systemic adaptation to changing physiological demands. All of the pancreatic endocrine cell types including the different beta-cell subpopulations originate from a common progenitor. Spatiotemporal environmental cues during development modify and restrict progenitor gene-expression profiles to drive differentiation and generate cellular heterogeneity. However, we have a limited understanding of the mechanisms that orchestrate differential gene expression patterns underlying *β*-cell heterogeneity.

Epigenetic mechanisms, such as DNA methylation, mediate context specific changes in gene-expression and direct the progressive refinement of cell fates during development (Avrahami and Kaestner 2012). DNA methylation is the most well studied epigenetic module, with the *de novo* DNA methyltransferases Dnmt3a and b establishing new DNA methylation patterns, while Dnmt1 maintains and propagates existing methylation patterns through cell division (Lyko 2018). Modulation of DNA methylation patterns has been implicated in the restriction of cell fate choices, and has been particularly well studied in neurogenesis (Parry et al. 2021). In pancreas, maintenance methylation by Dnmt1 is essential for the survival of pancreatic progenitors during embryonic development (Georgia et al. 2013), and for maintaining *β*-cell identity in postnatal life (Dhawan et al. 2011). Restriction of Dnmt1 expression also contributes to lineage commitment bias within the endocrine progenitor pool (Liu et al. 2019). Much less is known about the contribution of new methylation patterns in endocrine specification and cell-fate restriction. Dnmt3a is a part of the Nkx2.2 repressor complex that directs *β*-cell specification and is also essential for *β*-cell maturation in postnatal life (Papizan et al. 2011; Dhawan et al. 2015). This suggests that *de novo* DNA methylation is important in refining cell fate choices during differentiation.

Here, we focus on a unique *β*-cell subpopulation as a paradigm to illustrate the contribution of DNA methylation in guiding endocrine cell fate choices and heterogeneity. Pancreatic islets harbor a small number of cells that express the enzyme Tyrosine Hydroxylase (TH) which is a signature of sympathetic neurons; it catalyzes the rate-limiting step in the synthesis of catecholamines that inhibit insulin secretion (Teitelman et al. 1988; Morrow et al. 1993). The expression of TH in islets is developmentally regulated; it overlaps with glucagon in early pancreas development and later localizes to *β*-cells (Vazquez et al. 2014). Restriction of the TH+ sub-population of *β*-cells within islets is essential for appropriate insulin secretion in mice, such that mouse strains with larger number of TH co-expressing *β*-cells have blunted insulin secretion (Mitok et al. 2018). However, to date there have been no systematic studies characterizing the molecular and developmental identity of these cells, nor do we know the mechanisms that restrict TH expression in *β*-cells.

In this manuscript, we demonstrate the essential requirement of DNA methylation in establishing endocrine cell heterogeneity during differentiation, using TH expressing *β*-cells as a model. Our data establish that the islets cells co-expressing TH and insulin represent a *bona fide β*-cell subpopulation with distinct molecular and metabolic characteristics. Using developmental stage-specific ablation of DNA methyltransferases, we demonstrate that DNA methylation restricts TH expression in *β*-cells during the transition from endocrine progenitors to beta-cells. Loss of *de novo* DNA methyltransferase Dnmt3a in pancreatic- and endocrine-progenitor lineages results in a massive increase in TH-positive *β*-cells in postnatal life, while *β*-cell specific loss of Dnmt3a does not. Finally, we show that maintenance of TH promoter methylation is essential for the continued restriction of TH expression in postnatal *β*-cells, and loss of promoter methylation leads to increased TH expression and impairment of *β*-cell identity in response to chronic overnutrition. Overall, our data establish an essential role for DNA methylation patterning in endocrine cell fate determination and generation of cellular heterogeneity.

## Results

### The TH+ islet cells co-expressing insulin represents a *bona fide* β-cell subtype

Islet cells that co-express the enzyme TH have been described before (Teitelman et al. 1993), though their precise molecular identity has not been carefully defined. The TH expressing islet cells were readily detectable in adult wildtype mice, with majority of the islet TH+ cells co-expressing insulin (Fig. 1A-B, Supplemental Fig. S1A). These *β*-cells had variable levels of TH and followed a random distribution across islets, frequently appearing as clusters (Fig. 1B)., There was no overlap between TH and glucagon in adult islets, while TH was present in a rare few adult *δ*-cells (Supplemental Fig. S1B, C). TH expression in *β*-cells was first observed at embryonic day 15.5 (E15.5), prior to which TH was primarily restricted to *α*-cells (Supplemental Fig. S1D). To determine if this *β*-cell subpopulation exists in humans, we mined publicly available analysis of human islet single-cell RNA-seq (scRNA-seq) data (Mawla and Huising 2019), revealing that *TH* mRNA is expressed in very few *β*-cells in humans (Supplemental Fig. S1E). A survey of fetal, neonatal, and adult human pancreas did not show any TH+ cells in the limited number of samples analyzed (Supplemental Fig. S1F), confirming their rarity in humans. We confirmed the endocrine and *β*-cell lineage identities of the islet resident TH expressing cells by using endocrine (*Ngn3*- Cre:*R26R-mTmG*) and *β*-cell (*Ins1-*Cre:*R26R-YFP*) specific lineage reporters (Fig. 1C, Supplemental Fig. S1G), as well as by co-localization of TH with the endocrine marker Chromogranin A (ChrgA) (Fig. 1D). We asked if the TH+ *β*-cell represent a subpopulation with molecular features distinct from mature *β*-cells. TH+ *β*-cells expressed Glut2, *β*-cell transcription factors Pdx1, Nkx6.1, MafA, and NeuroD1, and maturity marker Ucn3 similar to other *β*-cells (Fig. 1E-G, Supplemental Fig. S1H-I). Given that TH also marks sympathetic neurons which are intimately linked to islet vasculature, we tested if the TH+ *β*-cells interacted with sympathetic neurons or vasculature (Reinert et al. 2014). The TH+ *β*-cells contacted the Tuj1+ sympathetic fibers as well as PECAM1+ capillaries (Fig. 1H, Supplemental Fig. S1J), suggesting that these cells mediate the interaction between islets and the islet neurovascular inputs. Prior work has shown an inverse correlation between the abundance of TH+ *β*-cells on islet function (Mitok et al. 2018). We tested the functional effect of inhibiting TH activity in mouse islets using the inhibitor AMPT (alpha-methyl-para-tyrosine), which led to improved glucose stimulated insulin secretion (GSIS) (Supplemental Fig. S1K), highlighting the functional significance of TH+ *β*-cells.

**Fig. 1.**
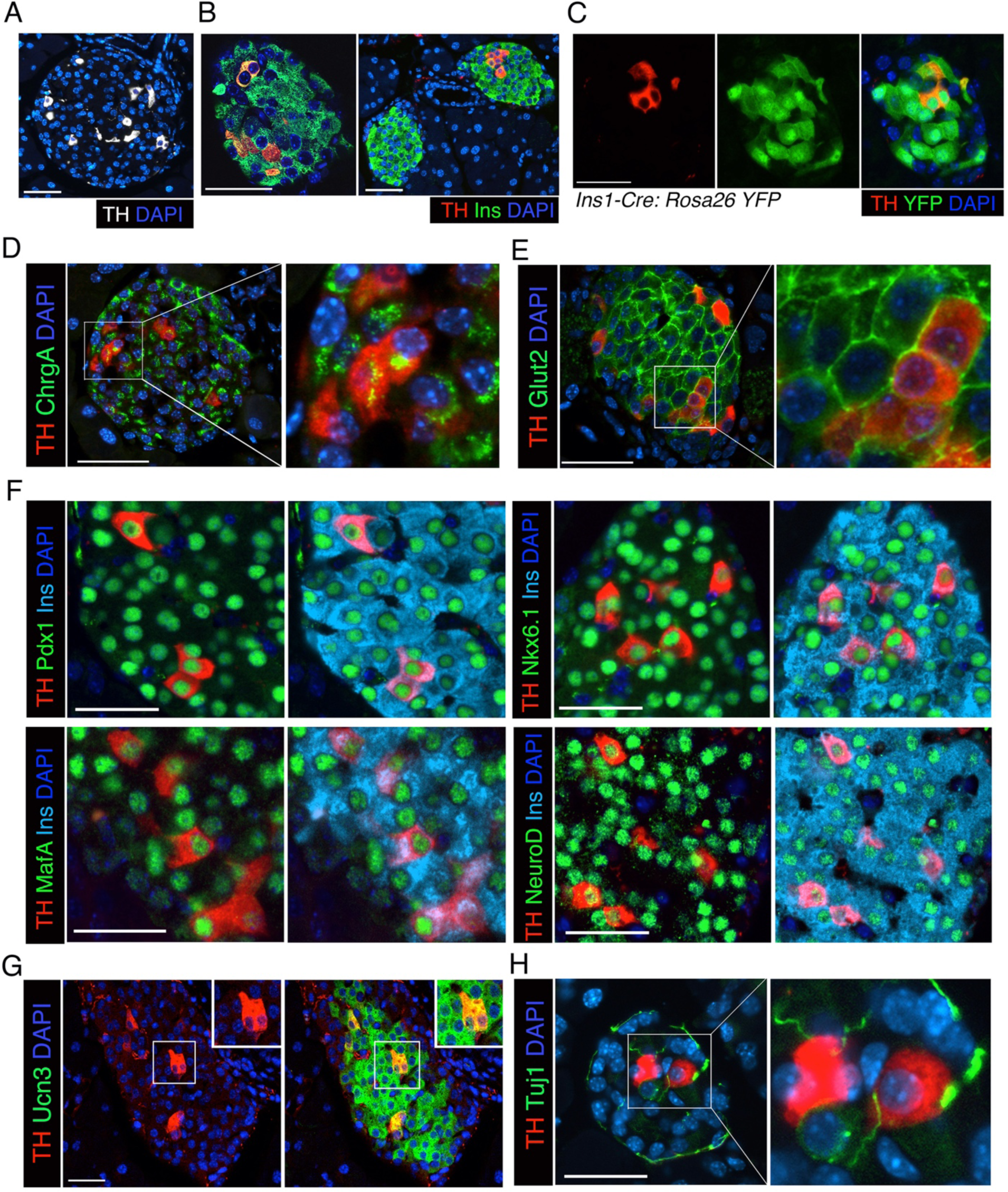
TH+ islets cells co-expressing insulin represent a *bona fide β*-cell subpopulation. *(A, B)* Immunostaining for Tyrosine Hydroxylase (TH: grey) *(A)*, TH and Insulin (TH: red; Ins: green) *(B)*, DAPI (blue). *(C)* Representative images from *Ins1*-Cre:*Rosa26* YFP mice (2 months) of age, stained for TH (red), YFP (green), DAPI (blue). *(D, E)* Immunostaining for TH (red) with chromogranin A (ChrgA : green) *(D)* or Glut2 (green) *(E),* with DAPI in blue. Right panels show a 2.5X magnified view of the area marked by white square. *(F)* Immunostaining for TH (red), Pdx1/Nkx6.1/MafA/NeuroD1 in green, with DAPI (blue) *(G)* Overlap of TH (red) with Urocortin3 (Ucn3: green) in adult (2 months old) wildtype pancreas. DAPI (blue). Inset shows a 2.5 X view of the area marked by white boxes. *(H)* Immunostaining for TH (red) with sympathetic neuronal marker Tuj1 (green). Right panel is a 2.5X magnification of the area in white box. *(B, D-F*) show representative images from adult (2 months) C57BL/6J mice, n=5 mice. Scale bar: 50 μm.

### TH+ β-cells in the adult pancreas are transcriptionally and metabolically distinct

To define the unique molecular features of TH+ *β*-cells, we meta-analyzed publicly available islet scRNA-seq data from adult C57BL/6J mice (Pineros et al. 2020). All samples were integrated into one dataset and various cell types were identified by their standard gene signatures cells (Fig. 2A, Supplemental Fig. S2A-B). *β*-cells were grouped into *Th*+ and *Th*- categories; 3 among the 4 *β*- cell clusters were enriched in *Th* (Fig. 2B). Next, we surveyed for differentially expressed genes (DEGs) between the *Th*+ and *Th*- *β*-cells and identified 130 such genes (Fig. 2C). The top enriched genes in the *Th*+ group were *Mt1*, *Ftl1*, *Selenow*, *Mt2*, *mt-Nd3*, while *Fkbp11*, *Manf*, *Pcsk1*, *Txnip*, *Pdia6, Ero1lb* represented top downregulated genes. Gene-set enrichment analysis (GSEA) of DEGs between *Th*+ and *Th*- *β*-cell states showed enrichment of pathways associated with protein synthesis, ribosomes, energy production, and ATP metabolism. The downregulated genes, on the other hand, were associated with endoplasmic reticulum (ER)- protein transport and ER stress response (Fig. 2D, Supplemental Fig. S2C). Collectively, this suggested that the TH+ *β*-cells may be predisposed to ER-stress and have higher oxidative phosphorylation (OxPhos).

**Fig. 2.**
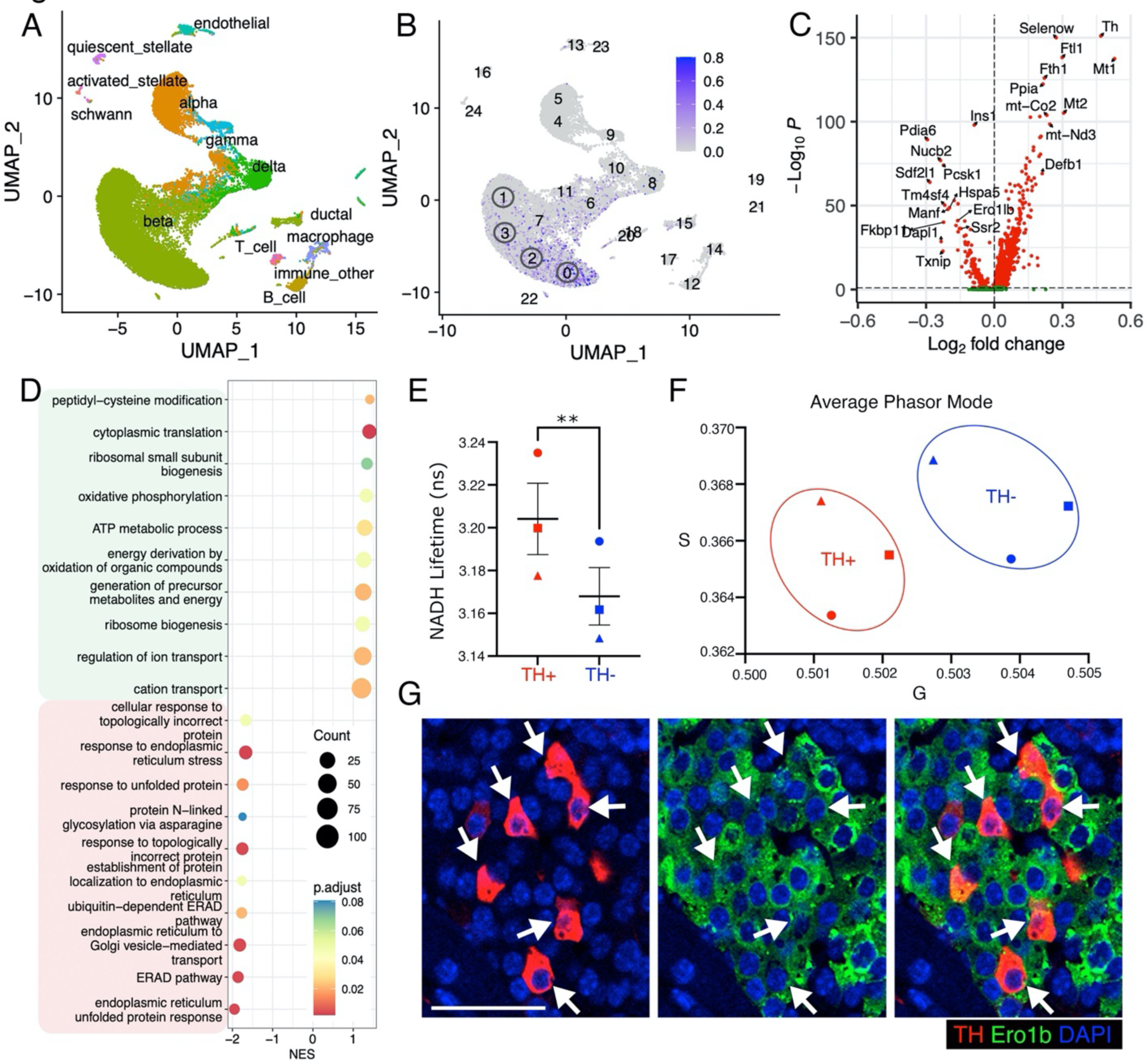
TH+ *β*-cells display distinct molecular and metabolic characteristics. *(A)* Analysis of scRNA-seq of mouse pancreatic islets from adult C57BL/6J mice on regular chow showing a Uniform Manifold Approximation and Projection (UMAP) plot with individual cell types marked with specific colors *(A)*. *(B)* Distribution of *Th* expression within *β*-cell subtypes. The four major *β*-cell clusters (0-3) are marked by open circles. *(C)* Volcano plot showing DEGs between *Th*+ vs. *Th-* cells in all *β*-cell clusters. *(D)* GSEA of the differentially expressed genes showing top enriched pathways for *Th*+ vs. *Th-* cells in *β*-cell clusters. The up- and down-regulated pathways are highlighted in green and red. Enrichment was calculated for Gene Ontology-Biological process (GO-BP) terms. *(E)* NADH lifetime signature, averaged and plotted for TH+ (red) vs TH- (blue) *β*-cells. *(F)* Lifetimes modes for single islets were transformed onto phasor plots and averaged for each individual pancreas. Modes of phasor plot for TH+ (red) vs. TH- (blue) for each individual animal (n=> 8 islets, 3 animals) were plotted onto a G vs. S graph (arbitrary units)*. (G)* Immunofluorescence for TH (red) and Erol1b (green), DAPI (blue) in pancreas from 2 months old C57BL/6J mice. Arrows mark TH+ cells. *(A-D)* is metanalysis of pooled scRNA-seq data from n=3 independent islet preparations from 9 weeks old C57BL/6J mice on regular diet. *(E, F)* Average of n=3 mice, => 8 islets/mouse. Error-bars show SEM. ns = not significant, ** *P*<0.01, *** *P*<0.005, using a paired *t*-test for *(E)*. *(G)* shows representative images from n=5 mice. Scale bar: 50 μm.

To validate these findings, we first determined the metabolic characteristic of the TH+ *β*-cells. For this, we leveraged a fluorescence lifetime imaging (FLIM) method developed by us to measure NADH levels in fixed tissue as a proxy of cellular metabolism (Millette et al. 2021). FLIM measures the lifetime of excited NADH; unbound NADH exhibits short lifetimes and is a byproduct of glycolysis; enzyme-bound NADH exhibits an ∼10X longer lifetime dependent on the bound enzyme and is a substrate of OxPhos. This allows FLIM to measure the glycolytic vs oxidative status of a cell that persists even after fixation (Lakowicz et al. 1992). We used immunofluorescence to identify TH+ and TH- *β*-cells and collected their fluorescence intensity and NADH auto-fluorescent lifetimes. Comparison of average lifetimes for TH+ vs TH- *β*-cells (n=>8 islets/pancreas; 3 mice) showed a higher lifetime for TH+ *β*-cells indicating more oxidative metabolism (Fig. 2E). We plotted the NADH intensity of each pixel of a TH+ and TH- *β*-cell in an islet onto a phasor plot and determined the coordinates of the mode of NADH intensity for individual islets. We then plotted the average of those modes of all islets per animal on a G vs. S axis (horizontal and vertical components of lifetime measurements in arbitrary units). The TH+ *β*- cells clustered together to the left, implying more OxPhos (Fig. 2F). Next, we tested the islet levels of the *Ero1lb* encoded Endoplasmic Reticulum Oxidoreductase 1 Beta (Ero1b) which catalyzes disulfide bond formation in the ER, is critical for insulin biogenesis, and protects from ER-stress. (Zito et al. 2010; Khoo et al. 2011). Immunostaining showed that Ero1b was less abundant in TH+ *β*-cells (Fig. 2G), confirming the scRNA-seq findings. Defective protein processing is a hallmark of the less differentiated *β*-cells (Lenghel et al. 2020; Shrestha et al. 2021), suggesting that TH+ *β*-cells may represent such a phenotype. To clarify this, we queried the “Beta-cell Hub” which features transcriptomic comparison of the immature and mature human *β*-cell states modeled by stem-cell derived *β*-like (sc-*β−*like) cells and human islet origin *β*-cells (Fang et al. 2019; Weng et al. 2020). Our query showed higher expression of *TH* in the sc-*β*-like cells compared to islet- derived *β*-cells, further indicating that TH marks a less differentiated *β*-cell state (Supplemental Fig. S2D). Collectively, the TH+ *β*-cells display signatures of poor protein processing and ER-stress response along with higher OxPhos, which are known to mark a less differentiated *β*-cell phenotype prone to ER-stress and oxidative stress (Newsholme et al. 2019; Lenghel et al. 2020).

### TH+ β-cells can replicate during postnatal life

It has been previously proposed that TH+ *β*-cells represent a senescent population which does not replicate (Teitelman et al. 1988). To determine whether TH+ *β*-cells are senescent, we tested the expression of senescence marker p21(Salama et al. 2014) in these cells. The TH+ *β*-cells in adult wildtype pancreas did not express p21, indicating that they are not senescent (Fig. 3A, Supplemental Fig. S3A). Replication is the primary mechanism for the growth and maintenance of *β*-cell mass in postnatal life, with rapid replication in neonatal life followed by gradual decline with age (Georgia and Bhushan 2004; Tschen et al. 2009). Therefore, we examined the growth and replication of TH+ *β*-cells at different stages in postnatal life. To establish the growth profile of TH+ *β*-cells, we quantified their percentage in pancreatic tissue from wildtype mice at various ages, ranging from postnatal day 2 (P2) to one year. We found that the percentage of TH+ *β*-cells increased steadily during the growth phase (P2-2.5 months), and then stabilized (Fig. 3B). Immunostaining of pancreatic sections from neonatal wildtype mice for TH and insulin along with replication markers Ki67 and pHH3 revealed that the TH+ *β*-cells do replicate in neonatal life (Fig. 3C-D). However, the number of replicating TH+ *β*-cells was highly variable compared to the majority non-TH+ *β*-cells, likely due to their small number in the islets (Fig. 3C-D, Supplemental Fig. S3B, C). In adult mice where *β*-cells are typically quiescent but can replicate in response to increased insulin demand (Dhawan et al. 2007), several TH+ *β*-cells were marked by the replication licensing factor Mcm2 (Blow and Hodgson 2002) indicating their replication competence (Supplemental Fig.S3D). Collectively, this shows that TH+ *β*-cells can replicate and are not senescent under homeostatic conditions.

**Figure 3.**
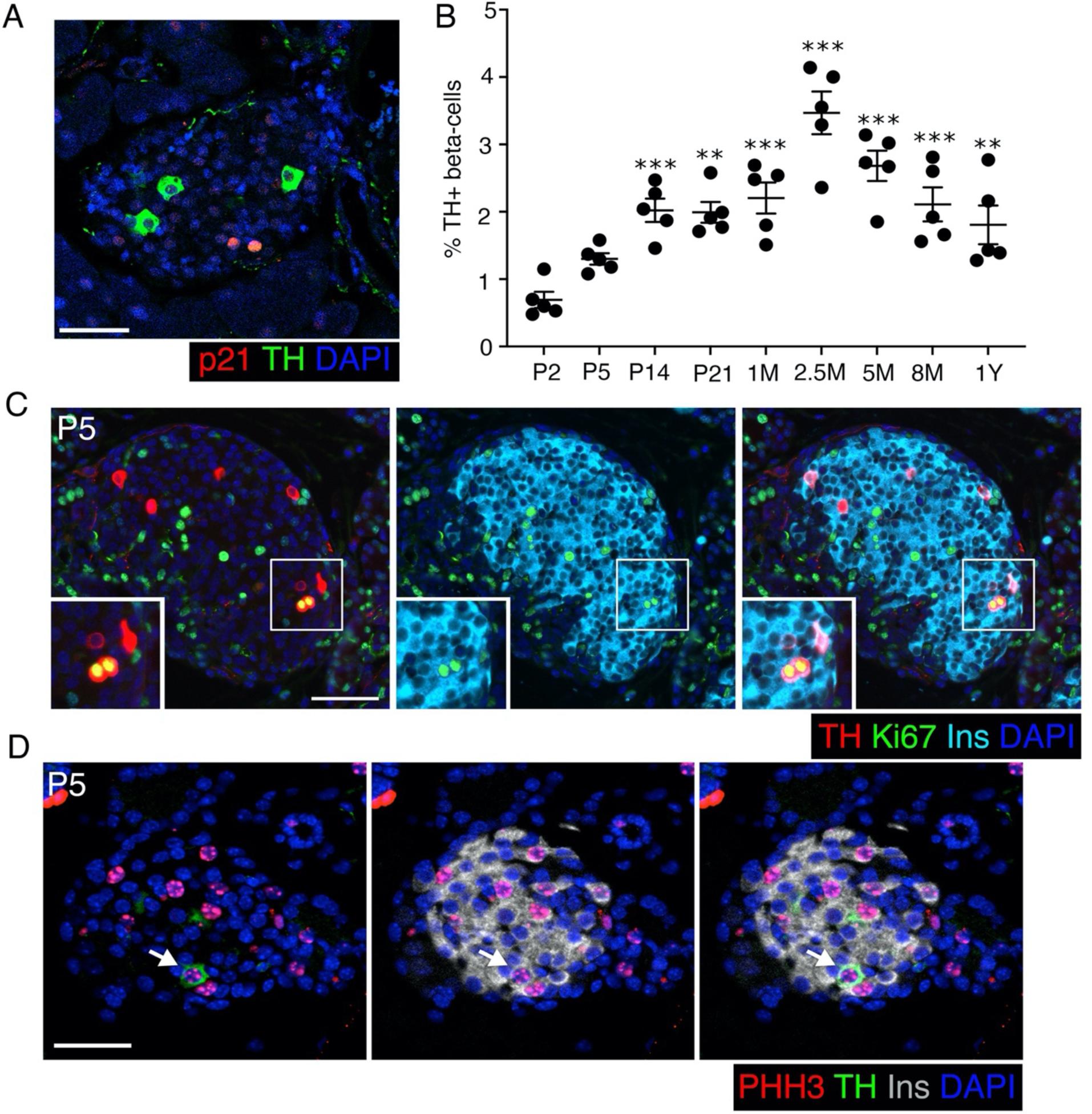
The TH+ *β*-cells can replicate and expand in postnatal life. *(A)* p21 (red) and TH (green) in pancreas from 2 months old mice. *(B)* Quantification of TH+ *β*-cells shown as percentage of total *β*-cells at indicated stages. *P* values shown are compared to P2. *(C)* Immunostaining for Ki67 (green), TH (red), insulin (cyan), with DAPI (blue) in P5 pancreas. Inset shows a 2X view of area marked by white boxes. *(D)* Phospho-histone H3 (pHH3: red) with insulin (grey), TH (green), DAPI (blue). All data is from wildtype C57BL/6J mice. Panels show representative or mean data from n=5 mice per group, with error-bars showing SEM of the mean. ** *P*<0.01, \*\*\**P*<0.005, using 1-way ANOVA with Bonferroni post-hoc test. Scale bar: 50 μm.

### TH promoter is methylated during the transition from endocrine progenitors to β-cells

DNA methylation patterning is a key mechanism that directs the establishment of cellular identity in pancreatic endocrine cells (Dhawan et al. 2011; Papizan et al. 2011). We therefore hypothesized that methylation of the TH promoter restricts the expression of TH to a select subset of *β*-cells in the islets. To test this, we first examined the DNA methylation at the promoter region of the *Th* locus in different stages of endocrine differentiation. We performed bisulfite sequencing for the *Th* promoter in pancreatic progenitors (Pdx1+), endocrine progenitors (Neurogenin3/Ngn3+), and *β*-cells (Insulin/Ins+) isolated from pancreatic tissue using lineage reporter mice at embryonic days 11.5 (*Pdx1*-Cre: *R26R-YFP*), 13.5 (*Ngn3*-Cre:*R26R-YFP*), and 18.5 (*Ins1-*Cre*^Thor^*: *R26R-YFP*), respectively. We found stage-specific, differential DNA methylation at the -2K region of the *Th* promoter, with the promoter being hypomethylated in the pancreatic and endocrine progenitors, and hypermethylated in sorted *β*-cells (Fig. 4A). No differences in DNA methylation were found in the other CG rich regions in the promoter, namely the -4K region and the proximal promoter region (Supplemental Fig. S4A-B). This suggested that the *Th* promoter is methylated during the transition from endocrine progenitors to *β*-cells.

**Figure 4.**
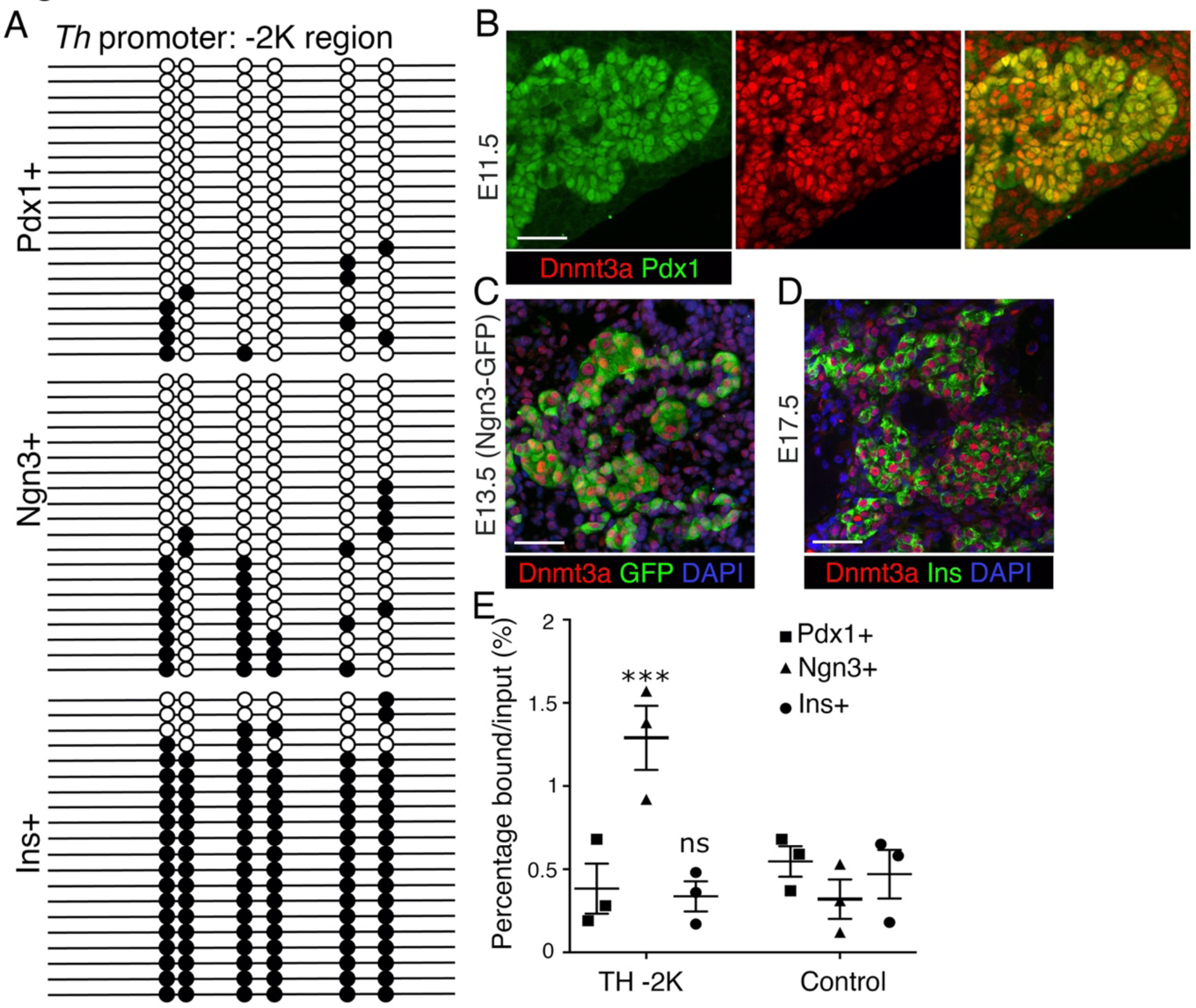
TH promoter undergoes Dnmt3a dependent methylation during endocrine progenitor to *β*-cell differentiation. *(A)* Bisulfite sequencing analysis of the *Th* promoter -2K region in purified mouse embryonic pancreatic progenitors, endocrine progenitors, and *β*-cells. Each line with dots is an independent clone; with filled and open circles denoting methylated and unmethylated CpGs, respectively. *(B-D)* Representative immunostaining for Dnmt3a (red); with Pdx1 (green) at E11.5 *(B)*, Ngn3-GFP (green) at E13.5 *(C)*, and insulin (green) at E17.5 *(D)*, (DAPI blue)*(C, D)*. (E) ChIP analysis for Dnmt3a binding to the -2K region of *Th* promoter, or a negative control region in sorted pancreatic progenitors, endocrine progenitors and TH+ *β*-cells. Panel *A* shows representative data from one of n=3 samples. Panel *E* shows the data as mean of n=3 independent samples per group, each sample being a pool of cells derived from multiple embryos, with error- bars showing SEM. \*\*\**P*<0.005, using 1-way ANOVA followed by a Bonferroni post-hoc test. Scale bar: 50 μm.

This differential methylation was not due to stage-specific expression of Dnmt3a (the only *de novo* Dnmt present in endocrine lineage, (Dhawan et al. 2015)), as Dnmt3a was expressed in the progenitors as well as *β*-cells (Fig. 4B-D). Chromatin immunoprecipitation (ChIP) for Dnmt3a recruitment at the -2K region of the *Th* promoter in the progenitors and *β*-cells revealed higher enrichment of Dnmt3a at the *Th* promoter in the endocrine progenitors, compared to pancreatic progenitors and *β*-cells (Fig. 4E). This further suggested that the methylation of *Th* promoter occurs during the transition from endocrine progenitors to *β*-cells. Expression analysis revealed very little overlap of Pdx1 and TH at E12.5 except a rare few TH+ cells marked by very low Pdx1, while none of the Ngn3+ endocrine progenitors expressed TH at E13.5 (Supplemental Fig. S4C- D). Given that TH is not expressed in the pancreatic- or endocrine-progenitors, the lack of promoter DNA methylation at these stages led us to determine the mechanism of *Th* repression at these stages. Histone H3-lysine 9 trimethylation (H3K9me3) is known to establish a reversible repressive chromatin state, prior to the establishment of the more stable DNA methylation (Lehnertz et al. 2003; Epsztejn-Litman et al. 2008). ChIP for H3K9me3 confirmed its enrichment at the -2K region of *Th* promoter in sorted pancreatic and endocrine progenitors (Supplemental Fig. S4E). These data show the stepwise establishment of repressive marks at the *Th* promoter through *β*-cell differentiation with DNA methylation as the final step, and suggest a potential role for DNA methylation in restricting TH expression in differentiated *β*-cells.

### Promoter DNA methylation is essential for the restriction of *Th* expression in β-cells

To establish if DNA methylation plays a direct regulatory role in restricting *Th* expression in *β*- cells during endocrine differentiation, we generated mice harboring loss of Dnmt3a in pancreatic, endocrine, and *β*-cell lineages and examined the TH expression pattern in pancreas. Mice with Dnmt3a ablation in the pancreatic and endocrine progenitor lineages (*Pdx1*-Cre:*Dnmt3a^fl/fl^*: termed 3aPancKO and *Ngn3*-Cre: *Dnmt3a^fl/fl^*: termed 3aEndoKO, respectively) showed a massive increase in the number of TH+ *β*-cells. Loss of Dnmt3a in the *β*-cell lineage (*Ins1-*Cre*^Thor^*: *Dnmt3a^fl/fl^*: termed 3aBetaKO) did not alter the number of TH+ *β*-cells (Fig. 5A-C, Supplemental Fig. S5A- C, G). We confirmed the efficiency of recombination and Dnmt3a ablation by Cre-driven fluorescent protein expression and Dnmt3a immunostaining. We chose neonatal stages to confirm Dnmt3a ablation as its levels decline dramatically post-weaning (Supplemental Fig. S5A-F) (Dhawan et al. 2015). Notably, loss of TH restriction upon Dnmt3a ablation appeared to be primarily in *β*-cells, suggesting lineage specificity of DNA methylation patterns during endocrine development (Supplemental Fig. S5G). In agreement with the changes in TH expression, the -2K region of the *Th* promoter was hypomethylated in islets isolated from 3aPancKO and 3aEndoKO mice compared to corresponding controls, while no significant difference was observed between islets from 3aBetaKO and controls (Fig. 5D-F). Collectively, our data show that *de novo* methylation by Dnmt3a is required to restrict *Th* expression in differentiated *β*-cells, and that temporally this restriction occurs during or after *β*-cell specification.

**Figure 5.**
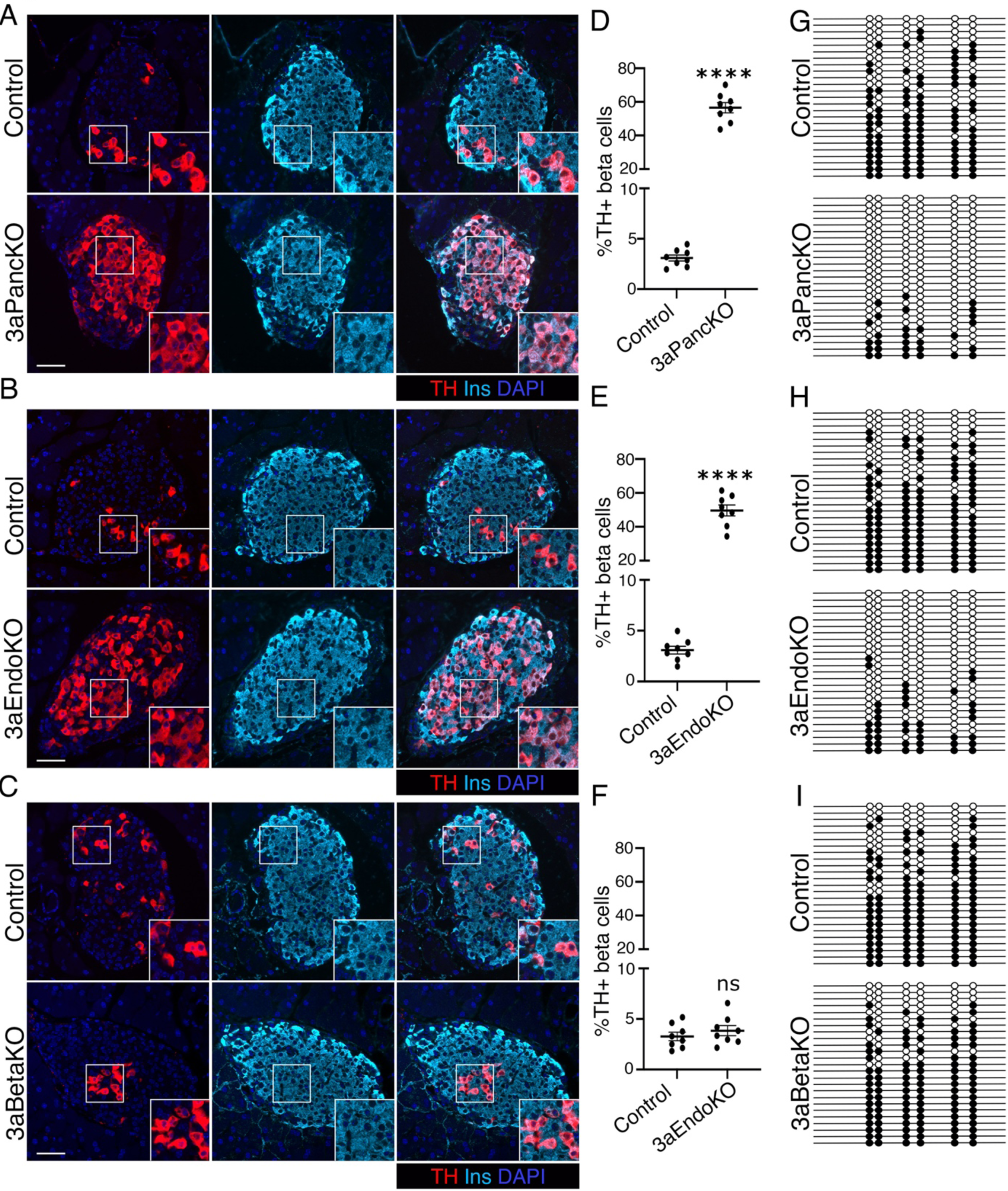
Dnmt3a ablation in progenitors but not *β*-cells leads to *Th* promoter demethylation and dysregulation in *β*-cells. *(A-F)* Representative images *(A-C)* and quantification of TH expression *(D-F)* in pancreatic sections from 2.5 months old mice with Dnmt3a ablation at various stages of pancreas development and littermate controls. TH (red), insulin (cyan), and DAPI (blue). Insets shows a 2X magnified view of areas marked by white boxes. *(G-I)* Bisulfite sequencing analysis of the *Th* promoter -2K region in islets from adult 2.5 months old 3aPancKO, 3aEndoKO, 3aBetaKO mice, and corresponding littermate controls. Each horizontal line with dots represents an independent clone and 25 clones are shown here; with filled circles representing a methylated CpG, while an open circle denotes an unmethylated CpG residue. Panels *(A-C)* show examples from n=5 independent samples, panels *(D-F*) show mean data from n=8 mice per group, with error- bars representing SEM.\*\*\**P*<0.005, determined by using a two-tailed Student’s *t*-test. Panels *G, H, I* show representative data for one of the n=5 independent islet preparations from single KO and control mice. Scale bar: 50 μm.

### DNA methylation dependent restriction of TH occurs prior to β-cell maturation and maintenance of these patterns is required for sustained TH repression

Our prior work showed the essential requirement for Dnmt3a dependent *de* novo DNA methylation in *β*-cell maturation (Dhawan et al. 2015). Therefore, we asked if the DNA methylation dependent restriction of TH expression in *β*-cells is linked to functional maturation. We compared TH+ *β*- cells in pancreatic sections from 3aPancKO and control littermates at postnatal day 0 and 5 (P0 and P5), before functional maturation. The increase in the number of TH+ *β*-cells in the 3aPancKO was very pronounced at P5, and evident even immediately after birth (Fig. 6A, E, Supplemental Fig. S6A-B). Dnmt3a ablation in the endocrine lineage showed a similar derepression of TH in *β*- cells at P0 (Figure 6B, F). To determine the developmental timeline of TH derepression in the absence of Dnmt3a, we examined 3aPancKO, 3aEndoKO, and control tissue during late endocrine differentiation (embryonic day 16.5: E16.5). We observed a clear increase in the number of TH+*β*-cells in the two KO models even at this stage when TH expression within *β*-cells is just beginning to emerge (Supplemental Fig. S6C, D). This suggests that DNA methylation restricts the expression of TH in *β*-cells during lineage specification, prior to their functional maturation.

**Figure 6.**
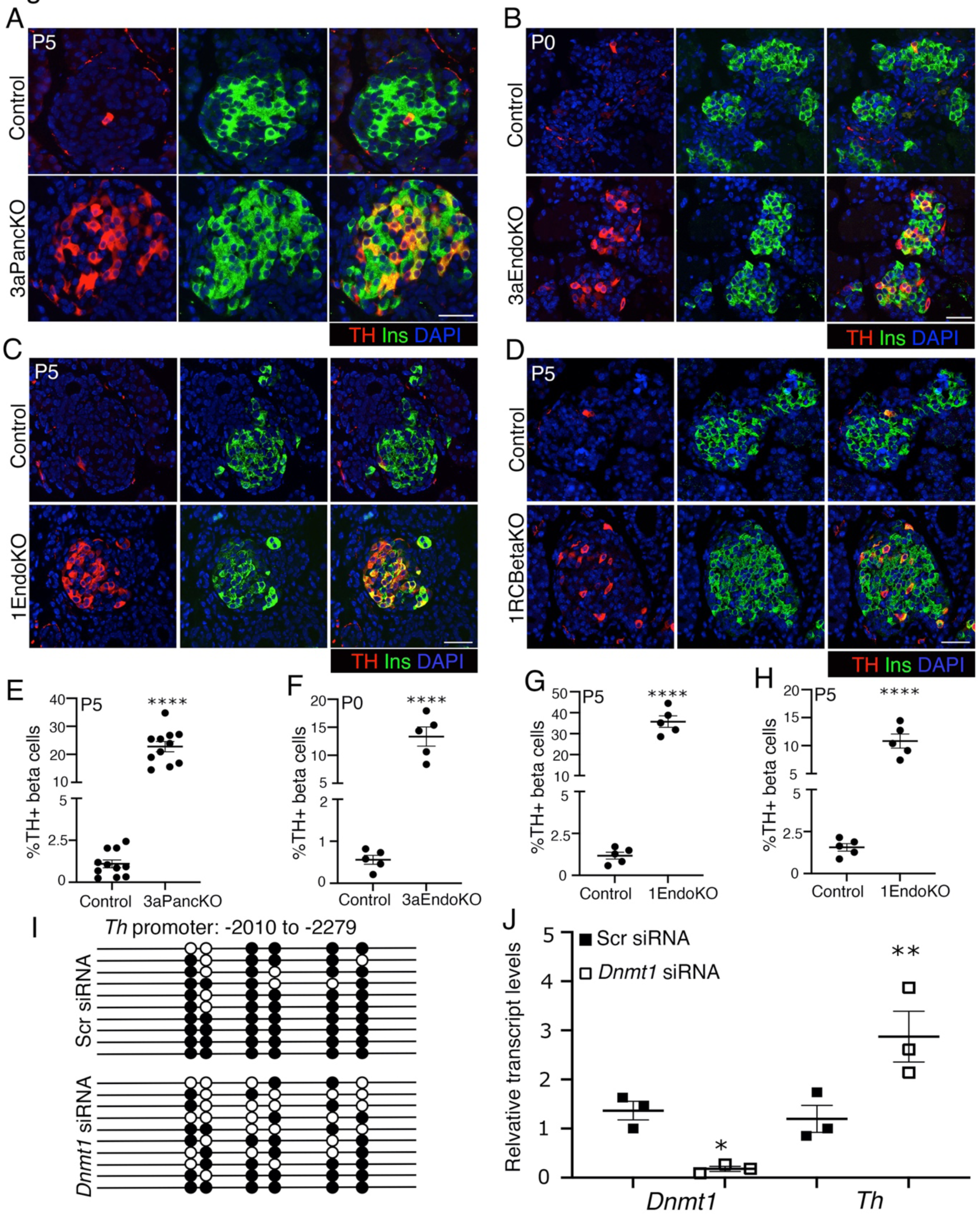
Dnmt3a dependent restriction of *Th* expression occurs prior to *β*-cell maturation and needs to be maintained for its continued restriction. *(A-H)* Immunofluorescence analysis *(A-D)* and quantification *(E-H)* of TH expression in pancreats from mice with Dnmt3a or Dnmt1 ablation in different lineages along with littermate controls at indicated ages. TH (red), insulin (green), and DAPI (blue). *(I-J)* Bisulfite sequencing of the -2K region of *Th* locus *(I)* and levels of *Dnmt1* and *Th* mRNAs relative to *Cyclophilin (J)*, in Min6 cells transfected with *Dnmt1* or scrambled (*Scr*) siRNAs. 10 independent bisulfite sequencing clones are shown in *(E)*; filled circles represent a methylated CpG, while open circles denote an unmethylated CpG residue. Data in *(A-H)* shows representative images or mean data from n=8 samples for 3aPancKO and controls, and n=5 for rest of the groups. Panel *I* shows representative data for one of the n=3 independent Min6 cell preparations treated with *Dnmt1* and Scr siRNAs, while panel *J* shows data from all n=3 preparation. Error-bars show SEM. \*\*\*\**P*<0.001, determined by using a two-tailed Student’s *t*- test for *(E-H)*, and using a 1-way ANOVA followed by Bonferroni’s post-hoc test for panel *J*. Scale bar: 50 μm.

Maintenance of the existing DNA methylation patterns through replication is essential for the preservation of *β*- vs *α*-cell identity in postnatal life (Dhawan et al. 2011). To determine if maintenance of *Th* promoter methylation is required for its sustained restriction in *β*-cells, we generated mice lacking the maintenance methyltransferase Dnmt1 in endocrine and *β*-cell lineages (*Ngn3*-Cre: *Dnmt1^fl/fl^*; 1EndoKO, and *RIP-*Cre*^Herr^*: *Dnmt1^fl/fl^*; 1RCBetaKO). As Dnmt1 depletion in *β*-cells results in their conversion to *α*-cells in adult life (Dhawan et al. 2011), we focused our analysis on neonatal life (P5) where the 1BetaKO *β*-cells have not yet trans-differentiated. We observed increased TH+ *β*-cells in the 1EndoKO and 1RCBetaKO mice at P5 (Fig. 6C-D), the increase being much higher in 1EndoKO congruent with the timeline of *Th* promoter methylation (Fig. 6G-H). To confirm whether failure to maintain DNA methylation directly contributes to *Th* upregulation upon *Dnmt1* ablation, we examined *Th* promoter methylation in a previously established model where synchronized Min6 cells treated with *Dnmt1* or scrambled (Scr) siRNA through three rounds of replication (8 days) to allow replication dependent DNA demethylation (Dhawan et al. 2011). Bisulfite sequencing showed loss of DNA methylation at the -2K region of the *Th* locus upon *Dnmt1* knockdown, corresponding to increased *Th* mRNA (Fig. 6I, J). Collectively, these data suggest that propagation of *Th* promoter methylation patterns is critical for maintaining the appropriate relative distribution of *β*-cell subpopulations.

### Promoter demethylation dysregulates TH expression in β-cells in response to long term high fat diet treatment

Given the essential requirement of maintenance DNA methylation in restricting TH-expression in *β*-cells, we asked if conditions that warrant *β*-cell expansion alter the proportion of TH+ *β*-cells. A prior study had reported increase TH+ *β*-cells in genetically obese *ob/ob* mice that undergo *β*- cell expansion in young age. However, the increase in TH+ *β*-cells only occurred in the older *ob/ob* mice (Teitelman et al. 1988). Such increase could either be age-related or due to chronically high insulin demand because insulin resistance. As wildtype mice do not show any age-dependent increase in TH+ *β*-cells (Fig. 3A), we hypothesized chronic increase in insulin demand to be the underlying cause. To test this, young adult wildtype mice (7 weeks old) were either exposed to a short-term (8 weeks) or long-term (16 weeks) high-fat diet (HFD) regimen, or maintained on a control diet (CD). The two regimens are designed to model successful *β*-cell adaptation to increased demand for insulin (short-term HFD) or the subsequent decompensation as *β*-cell de- differentiation and failure ensues (Weir and Bonner-Weir 2004). As expected, both short- and long-term HFD led to increased body weight and impaired glucose tolerance (Supplemental Fig. S7A-D). Analysis of the pancreatic tissue at end-point showed an increase in islet size and *β*-cell mass in mice on both short- and long-term HFD. However, only the long-term HFD regimen resulted in an increase in TH+ *β*-cells (Fig. 7A-F). This suggested that chronic *β*-cell workload due to persistently high insulin demand could disrupt the epigenetic restriction of TH in *β*-cells. To establish this, we performed bisulfite-sequencing analysis of *Th* promoter in islets from mice treated with short- and long-term HFD vs appropriate controls. Our data showed loss of promoter DNA methylation corresponding increase in *Th* mRNA expression in islets from mice on long-term but not short-term HFD (Fig. 7F, Supplemental Fig. S7 E F), indeed suggesting a failure to maintain DNA methylation patterns under conditions of chronic insulin demand. Collectively, these data suggest that maintenance of epigenetic patterns is essential to preserve *β*-cell identity during *β*-cell adaptation, and that such mechanisms collapse during *β*-cell decompensation.

**Figure 7.**
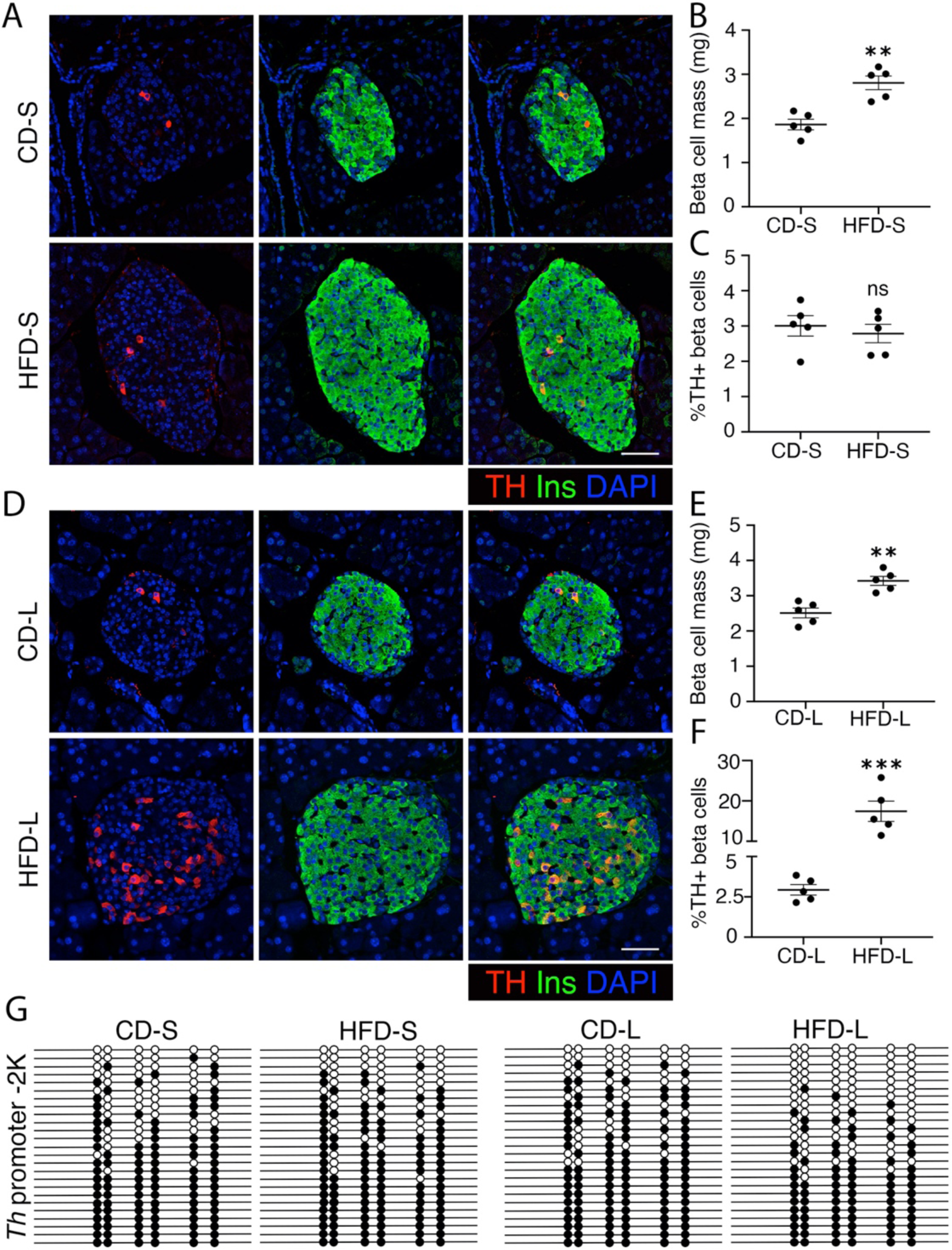
Chronic high fat diet leads to promoter demethylation dependent dysregulation of *Th* expression in *β*-cells. *(A-C)* Immunofluorescence for TH (red) and insulin (green), with DAPI in blue *(A)*, *β*-cell mass *(B)*, and quantification of TH+ *β*-cells *(C)* in wildtype 7 weeks old mice on a high fat- or control-diet for a short term (HFD-S or CD-S). *(D-F)* Immunostaining for TH (red) insulin (green) and DAPI (blue) *(D)*, *β*-cell mass *(E)*, and quantification of TH-positive beta-cells *(F)*, in wildtype 7 weeks old mice on high fat- or control-diet for a long term (HFD-L or CD-L). *(G)* Bisulfite sequencing analysis of the *Th* promoter -2K region in islets from mice fed with HFD or CD for 8 weeks (HFD-S, CD-S) or 16 weeks (HFD-L, CD-L. Each horizontal line with dots represents an independent clone and 25 clones are shown for each sample; with filled circles= methylated CpG, open circles unmethylated CpG. Immunofluorescence data shows representative images from n=5 samples. The data on *β*-cell mass and TH+ *β*-cell quantification shows mean of n=5 independent samples per group, while the bisulfite-sequencing data shows representative clones from one of the n=3 independent samples per group. Error-bars show SEM of the mean. Ns=not significant, \*\**P*<0.01, \*\*\**P*<0.005, determined by using a two-tailed Student’s *t*-test. Scale bar: 50 μm.

## Discussion

Pancreatic *β*-cells share several features with neurons including shared developmental transcriptional programs (Eberhard 2013). Islet cells are known to contain neurotransmitters that regulate insulin secretion and enzymes such as TH that regulate their synthesis (Teitelman and Lee 1987; Iturriza and Thibault 1993; Rodriguez-Diaz et al. 2011). We observe TH expression in a small subpopulation of adult *β*-cells and rare *δ*-cells, suggesting that neuronal-like features underlie heterogeneity of many endocrine cell types. Early endocrine precursors express specific neuronal markers, whose expression is restricted as endocrine identities are refined during development. Accordingly, it has been postulated that the TH+ *β*-cells indicate a developmentally immature precursor in the adult pancreas (Teitelman and Lee 1987; Rodnoi et al. 2017; Veres et al. 2019). Our data suggest that while the TH+ *β*-cells harbor many mature *β*-cell markers, they appear to represent a less differentiated state. The apparent lack of TH+ endocrine cells in developing and adult human pancreas along with abundant *TH* expression in sc-*β*-cells not only highlights species differences, but also points to differences between human endocrine differentiation *in vivo* and *in vitro*. Recent studies have similarly identified *β*-cells harboring serotonergic markers in human stem-to-*β*-cell differentiation (Veres et al. 2019). This suggests that expression of “neuronal-like” features characterize sc-*β*-cells, revealing a bottleneck in generating mature human sc-*β*-cells.

Our work shows that DNA methylation patterning is essential for the control of endocrine identity and heterogeneity in homeostatic and stress conditions. The specification and maintenance of cell-fates requires the precise temporal orchestration of epigenetic patterns to regulate stage-specific transcriptional programs (Golson and Kaestner 2017). *De novo* DNA methylation is essential for the repression of *α*-cell fate determinants in *β*-cells to guide *β*-cell lineage commitment in fetal life, and for *β*-cell maturation in early postnatal life. Maintenance of these methylation patterns by Dnmt1 through replication allows the continued restriction of *α*-cell fate to preserve *β*-cell identity in postnatal life (Dhawan et al. 2011; Papizan et al. 2011; Dhawan et al. 2015). We now show that the *de novo* DNA methylation dependent restriction of TH expression in *β*-cells occurs during lineage specification prior to their functional maturation. Heterogeneity of Dnmt expression within the endocrine progenitor pool dictates their lineage commitment bias (Liu et al. 2019); similar mechanisms might be involved in limiting TH expression to a select few *β*-cells. The ablation of *Dnmt1* in endocrine progenitors and *β*-cells results in the ectopic TH expression in *β*-cells prior to their trans-differentiation to *α*-cells. These temporal changes in *β*-cell identity upon the loss of maintenance methylation are intriguing and suggest differences in the stringency of restriction between cell-fates with different degrees of relatedness. It is likely that the epigenetic barriers between different *β*-cell subtypes are more plastic, compared to those between *β*- and *α*-cells. DNA methylation plays distinct regulatory roles in different pancreatic endocrine lineages; while loss of Dnmt1 in *β*-cells results in their conversion to *α*-cells, Dnmt1 ablation in *α*- or *δ*-cells does not convert them to *β*-cells (Dhawan et al. 2011; Chakravarthy et al. 2017; Damond et al. 2017). This points to high degree of specificity in DNA methylation dependent control of cell fates.

It has been suggested that TH+ islet cells represent a senescent population (Teitelman et al. 1988). We show that TH+ *β*-cells do replicate during the neonatal growth phase and do not harbor markers of senescence under homeostatic conditions. Instead, these cells bear signatures of over-active protein synthesis and poor ER-stress response, and higher oxidative phosphorylation, which are known hallmarks of a *β*-cell phenotype that is less differentiated and predisposed to ER-stress and oxidative stress (Newsholme et al. 2019; Lenghel et al. 2020). We also show that conditions of *β*- cell decompensation can disrupt the epigenetic repression of the *Th* promoter and alter *β*-cell identity marked by an increase in TH+ *β*-cells. These data explain prior work showing increased TH+ *β*-cells in older genetically obese *ob/ob* mice without any accompanying replication (Teitelman et al. 1988). The older *ob/ob* mice represent a *β*-cell decompensation state in response to chronically elevated insulin demand, where *β*-cell replication does not occur (Weir and Bonner-Weir 2004). The accumulation of these cells during *β*-cell decompensation, along with the inverse correlation between the abundance of TH+ *β*-cells and islet insulin secretion capacity (Mitok et al. 2018), suggests the possibility of *β*-cells de-differentiating into a TH+ state and acquiring a senescent, dysfunctional phenotype. Indeed, chronic *β*-cell stress is known to induce de- differentiation and senescence (Talchai et al. 2012; Thompson et al. 2019). The large scale changes in human islet methylome associated with *β*-cell identity and function genes in type 2 diabetes strengthen this idea (Volkmar et al. 2012). Our data suggest the relevance of early developmental DNA methylation patterning in the postnatal *β*-cell identity. This is significant, given that adverse exposures such as over- or under-nutrition during development can predispose to future risk of *β*- cell failure (Parveen and Dhawan 2021). Future work will identify what specific aspects of mature *β*-cell identity rely upon early developmental epigenetic patterning, and clarify the significance of such a regulatory paradigm.

## Materials and Methods

### Human pancreatic samples

Pancreatic sections from human late fetal and early neonatal pancreas (N=1 each) and adult non-diabetic donors (N=3) were procured from brain-dead organ donors by the JDRF Network for Pancreatic Organ Donors with Diabetes (nPOD), University of Florida at Gainesville (Campbell-Thompson et al. 2012). Human fetal pancreas (N=1; 16 weeks pc) was obtained through the Terminal Tissue Bank at the University of Southern California. 4 μm sections of pancreas from each subject were provided to the investigators, coded to conceal the personal identity of the subjects. The sample characteristics are described in Supplemental Table 1.

### Animal models

All animal experiments were performed in accordance with the National Research Council Guidelines for the Care and Use of Laboratory Animals, and under protocols approved by the Institutional Animal Care and Use Committee (IACUC) at City of Hope and USC, as well as the lead contact’s prior institution (UCLA). Mice were maintained by mating males and females on a C57BL/6J background. DNA methyltransferase Dnmt3a and Dnmt1 were ablated in different pancreatic lineages using the Cre-lox system. *Dnmt3a^fl/fl^* and *Dnmt1^fl/fl^* mice have been described previously (Jackson-Grusby et al. 2001; Kaneda et al. 2004). We used *Pdx1*-Cre (Hingorani et al. 2003) and *Ngn3*-Cre (Schonhoff et al. 2004) mice to ablate Dnmt3a or Dnmt1 in the pancreatic-and endocrine-progenitor lineages respectively, while the *Ins1-*Cre*^Thor^* (Thorens et al. 2015) or *RIP*-Cre*^Herr^* (Herrera 2000) were used for *β*-cell-specific deletion. The Cre lines also harbored the Rosa26R-*YFP*, or Rosa26R-mTmG lineage reporters (Soriano 1999). We used the *Pdx1*-Cre: Rosa26R-*YFP* embryos to sort pancreatic progenitors, while endocrine progenitors were harvested from the heterozygous Ngn3-EGFP mouse embryos (Lee et al. 2002). Transgenic mice expressing mouse *Ins1* promoter (*MIP-GFP*; (Hara et al. 2003)) were used to sort *β*-cells. Mice were fed *ad libitum* and kept under a 12hr. light/dark cycle. Both male and female mice were used for all studies, except the high fat diet (HFD) studies. For the HFD studies, 1.5 months old male C57BL/6J mice (n=8/group) were fed with either control diet (4.4% calories from fat; Envigo), or HFD (55% calories from fat, Envigo TD.93075) for short-term (8 weeks) or long-term (16 weeks).

### Immunostaining, imaging, and morphometric analyses

Standard immunofluorescence protocol was used for the detection of various proteins in pancreatic sections as previously described (Tschen et al. 2009; Rodnoi et al. 2017). Primary antibodies were diluted in the blocking solution as noted in Supplemental Table 2. Donkey- and goat-derived secondary antibodies conjugated to FITC, Cy3, Cy5, Alexa 448, 592, and 647 were diluted 1:200 (Jackson ImmunoResearch). An antifade mounting medium with DAPI was used to label nuclei (Vector Labs). Slides were viewed using a Leica DM6000 or DM6B microscope (Leica Microsystems) and images acquired using OpenLab (Improvision) or Leica Application Suite X (LAS X, Leica). Confocal imaging was done on the Zeiss LSM 800 platform with AiryScan using the Zeiss Efficient Navigation (ZEN) software. To quantify TH+cells, all islets were imaged and the total number of insulin+, TH+, and TH+Insulin+ cells was manually counted, the data was expressed either as percentage of total TH+ cells (Supplemental Fig. S1C), or *β*-cells and islet cells(Fig. 3A, B.) Ki67, pHH3, and Mcm2 were used as replication markers, and quantification of replication in different subpopulations was done as described in (Rodnoi et al. 2017). *β*-cell mass was measured per published protocols (Tschen et al. 2009).

Fluorescence Lifetime Imaging Microscopy (FLIM) was performed on SP8 DIVE FALCON multi-photon FLIM microscope (Leica Microsystems) using 40x/1.10 N.A. water immersion objective. NAD(P)H was excited with a Spectra-Physics Insight 3X ultrafast IR laser at 740 nm, 0.8mW average power, and 4 frame accumulations per optical section, 440-500nm emission. The Alexa dyes were excited using an 860 nm wavelength with the same Spectra-Physics laser. Images were taken at 1024 × 1024 resolution for N= 3 biological samples and >8 islets/sample. To analyze the metabolic signatures for each *β*-cell type, masks for regions of *β*-cells were created from fluorescent images of TH and insulin staining respectively by thresholding. Masks (TH+INS+ and TH-INS+) were then preprocessed to fill the cytoplasm of cells and exclude the nuclei. Each mask was applied to the field of view to extract lifetime information from TH+ and TH- cells separately. Paired *t*-test was used to determine significant differences between the two populations.

### Islets and cell lines, transfection, cell sorting

Islets were isolated from mouse pancreas using perfusion with the Liberase enzyme blend (Sigma Aldrich) as described (Tschen et al. 2009). Min6 cells were maintained as described previously in DMEM containing 10%FBS and 25 mM glucose at 37 °C in 5% CO2 environment. Knockdown of *Dnmt1* in Min6 was performed by transfection with a pool of specific targeting small inhibitory RNAs on-TARGETplus (siRNAs), or scrambled controls (Dharmacon Horizon Discovery). Transfections were performed in OPTI-MEM medium using Lipofectamine-RNAiMAX (ThermoFisher), according to manufacturer’s instructions. Cells were synchronized by serum starvation O/N, transfected with *Dnmt1* or scrambled siRNAs and harvested 8 days post-transfection for subsequent applications to allow for replication dependent DNA demethylation (Dhawan et al. 2011). For cell sorting, pancreatic buds from *Pdx1*-Cre: Rosa26R-*YFP* embryos, or heterozygous Ngn3-EGFP embryos, or the MIP-GFP transgenic mice were harvested, pooled, and digested into single cells using TrypLE Express (ThermoFisher). Cells were sorted for GFP using BD FACS Aria II to an average percentage purity of 85-95%, with wildtype cells as negative control for FACS gating (Dhawan et al. 2015).

### Metabolic and physiological analyses

Intra-peritoneal glucose tolerance tests (IP-GTT) and static incubation GSIS were performed as previously described (Tschen et al. 2009; Dhawan et al. 2015; Rodnoi et al. 2017). For TH inhibition, freshly isolated islets were cultured for 24 hours in the presence of 10 μM AMPT or vehicle (1X PBS) and used for GSIS. Insulin in supernatants and islet extracts was measured by ELISA (Mouse Insulin ELISA, Mercodia).

### ChIP, DNA methylation, and mRNA expression analyses

ChIP on the sorted progenitors and *β*-cells was carried out according to a low-cell number micro- ChIP protocol, with minor modifications (Dahl and Collas 2008; Dhawan et al. 2015). The antibodies and primers used for ChIP are listed in the Supplemental Tables 3 and 4. Bisulfite conversion of DNA from sorted cells and islets and sequencing of the converted DNA was performed according to established methods (Millar et al. 2002; Dhawan et al. 2011) using primers described in Supplemental Table 5. We analyzed 20-25 independent clones per group, each experiment repeated in biological triplicates. RNA extraction and quantitative Real-time RT-PCR was performed as published (Rodnoi et al. 2017), using primers listed in Supplemental Table 6.

### scRNA sequencing analysis

The analysis was performed on publicly available scRNA-seq datasets (n=3) from islets of mice fed with low fat diet (Piñeros et al. 2020), accession number GSE12512. The datasets were subjected to quality control (elimination of cells with RNA features < 200 or > 10,000 and mitochondrial RNA > 15%) and normalization (“LogNormalize” with scale factor 10,000) in Seurat 4.1 (Hao et al. 2021) retaining the top 2000 variable genes, followed by integration using the Seurat workflow (Stuart et al. 2019). The integrated data was scaled and subjected to principal component analysis (PCA), following by clustering using the top 15 PCs. The cell types were assigned via the SingleR package (Aran et al. 2019), using annotated data from mouse pancreatic cells as reference (Baron et al. 2016). The identified cell types were validated by comparing the expressions of known cell type specific biomarkers. The *β*-cells were further filtered for low expressions of *α*- and *δ*-cell markers. *β*-cells with *Th* expression > 0 were classified as *Th*+. The differential gene expressions were determined using the FindMarkers function in Seurat. For Genset enrichment analysis (GSEA), the differentially expressed genes were ranked according to their adjusted *p*-values, with signs based on the direction of the fold change. GSEA were performed for the Gene Ontology biological process terms (Ashburner et al. 2000; Carlson 2019; Collaborators 2021) using the R package ClusterProfiler (Yu et al. 2012). The enrichment and pathway module plots were prepared using the DOSE (Yu et al. 2014) and the enrichplot (Yu 2022) R packages respectively. The volcano plots were prepared using the R package EnhancedVolcano (Blighe et al. 2021).

### Statistical methods

All data were expressed as Mean±S.E. (standard error). Mean and SEM values were calculated from at least triplicates of a representative experiment. The statistical significance of differences was measured by unpaired Student’s *t*-test for experiments with two groups and a continuous outcome, while a one-way ANOVA with Šídák or Bonferroni post-hoc tests was used for experiments with repeat measures. For differences in blood glucose levels during the IPGTT, a repeated two-way ANOVA (time and treatment) was performed and significance determined using Bonferroni’s *post hoc* test. A *P* value < 0.05 indicated statistical significance. \**P*<0.05, \*\**P*<0.01, \*\*\**P*<0.005.

### Competing interest statement

The authors have declared that no conflict of interest exists.

## Acknowledgements

We thank Alexander Ham and Pope Rodnoi (Dhawan Lab) for technical support. We would like to thank Dr. Rachel Steward and Family Planning Associates for the (fetal) tissue donation (CHLA tissue repository) and Drs. Jason Junge and Peiyu Wang (USC) for expertise and consultation in FLIM studies. We thank Dr. David Ron (University of Cambridge) and Dr. Mark Huising (University of California, Davis) for the kind gifts of Ero1b and Ucn3 antibodies, respectively. We acknowledge funding support from the NIH (R01DK120523 to S.D., R01DK119590 to H.S.), Human Islet Research Network (a New Investigator Award to S.D. via UC4DK104162), The Homer & Gloria Harvey Family Foundation, The Paul Lester Foundation, and The Saban Research Institute (to S.K.G.). Work in the S.D. and H.S. laboratories is also supported by City of Hope (start-up support to S.D. and H.S.), Wanek Family Foundation to Cure Type 1 Diabetes (S.D. and H. S.), and Lions Club Initiative (S.D.).

## Author contributions

S.D. and S.K.G. conceived and planned the study. J.K.W., N.P., M.S.R., J.C., S.K.G. and S.D. performed the experiments. J.K.W., N.P., M.S.R., J.C., S.B., X.W., S.K.G., and S.D. performed the analyses. S.D., S.K.G., S.B., X.W., H.S., J.C., J.K.W., and N.P. interpreted data. N.P., S.B., X.W., S.K.G., H.S., and S.D. wrote the manuscript. S.K.G., H.S., and S.D. acquired funding.

---

**Supplemental Fig. S1.**
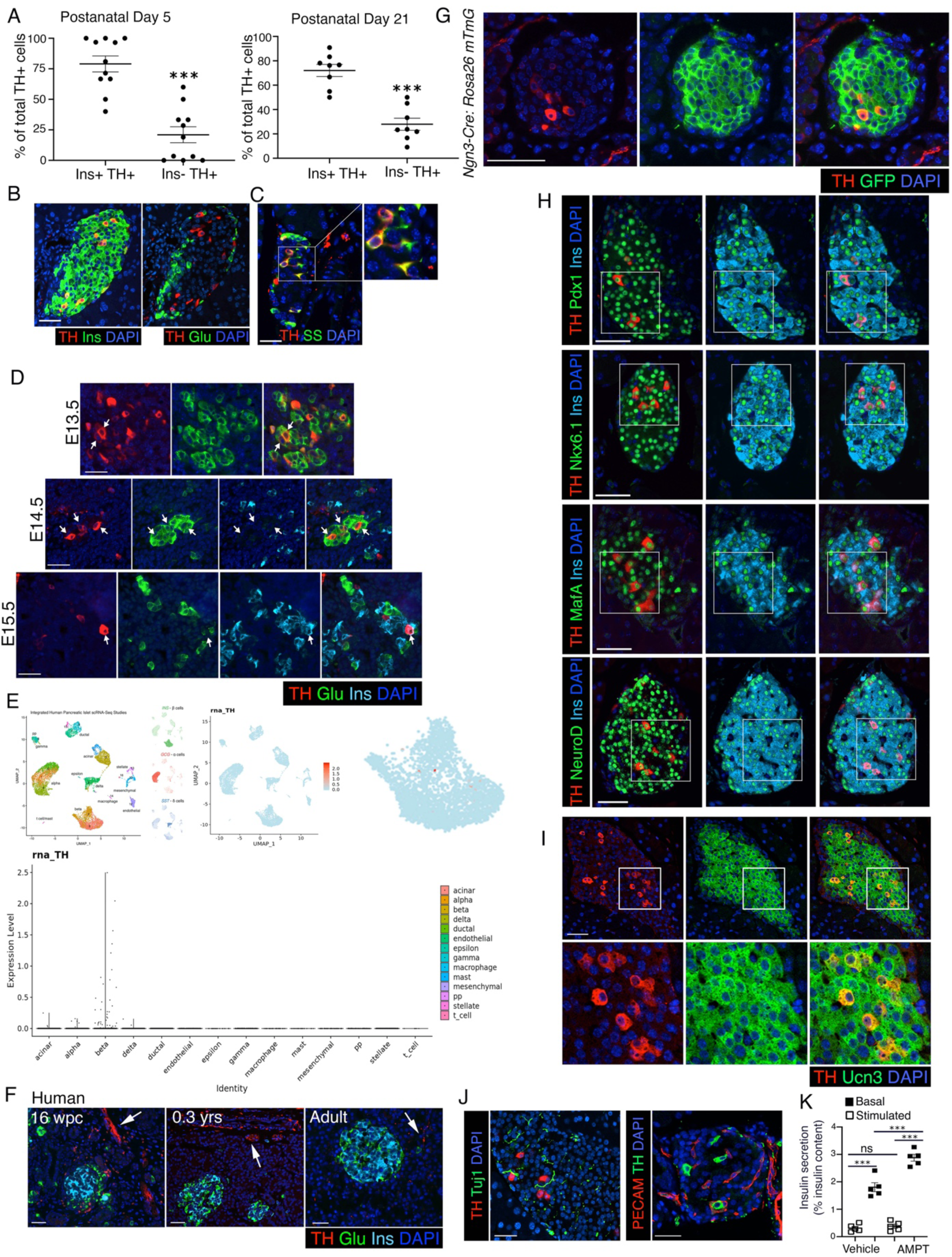
(related to **Fig 1).** *(A)* Immunofluorescence analysis of adjacent pancreatic sections from adult wildtype mice (2 months old) for TH (red) and insulin (Ins: green) in the left panel, and TH (red) and glucagon (Glu: green) in the right panel, showing that TH overlaps with insulin but not glucagon in the adult islets. DAPI counter-stains the nuclei (blue). *(B)* Immunostaining for TH (red) and somatostatin (SS: green), with DAPI (blue), showing that TH also marks a fraction of adult delta cells. Right panel shows a 2X magnified view of the area marked by a white square. *(C)* Morphometric quantification of TH-expressing cells assessed for insulin expression (shown as percentage of total TH-positive cells) at postnatal days 5 (left panel, n=11 wildtype mice) and 21 (right panel, n=8 wildtype mice), showing that majority of TH- expressing cells in the postnatal pancreas are beta cells. *(D)* Immunostaining for Tyrosine Hydroxylase (TH: red), glucagon (Glu: green), and Insulin (Ins: cyan), with DAPI in blue in embryonic pancreas at E13.5, E14.5, and E15.5. Arrows indicate cells co-expressing TH and Glu (top and middle panels; E13.5 and E14.5) or TH and Ins (bottom panel; E15.5). *(E)* Data shows distribution of *TH* mRNA expression in different endocrine cell populations in single-cell RNA sequencing (scRNA-seq) data meta-analyzed and made publicly accessible by (Mawla and Huising 2019) https://www.huisinglab.com/diabetes_2019/index.html. The data shown is an integration of four human pancreatic islet datasets, as noted in the study. Top panel shows a UMAP (Uniform Manifold Approximation and Projection) plot, while bottom panel shows a violin plot. *(F)* Immunostaining for TH (red), glucagon (Glu; green), and insulin (Ins; cyan), with DAPI (blue) in fetal (16 weeks p.c./gestation), neonatal (0.3 years) and adult human pancreatic sections, showing absence of TH in human islets. Arrows mark sympathetic nerve fibers that express TH, serving as an internal positive control. *(G)* Representative pancreatic sections from *Ngn3*-Cre:*Rosa26* mTmG lineage reporter mice at 2 months of age, stained for TH (red) and GFP (green) with DAPI in blue. *(H)* Full, un-cropped images corresponding to Fig. 1*(G)* for immunofluorescence analyses showing overlap of TH (red) with key transcription factor hallmarks of beta cell identity, namely, Pdx1 (top left), Nkx6.1 (top right), MafA (bottom left) and NeuroD1 (bottom right) shown in green. Nuclei are marked in blue with DAPI. The regions marked by white box are zoomed 2X and presented in Fig. 1*(G)*. *(I)* Overlap of TH (red) with mature beta cell marker Urocortin3 (Ucn3: green) in adult (2 months old) wildtype pancreas. DAPI (blue) counterstains the nuclei. Inset shows a 2.5X zoomed view of the area marked by white boxes. *(J)* Immunostaining for TH (red) with sympathetic neuronal marker (shown in green) Tuj1 (left) and Nestin (right), showing interaction of sympathetic afferents with TH-positive islet cells. *(K)* Static incubation GSIS in islets from 8 weeks old C57BL/6J mice, pretreated either with 10 μM AMPT or vehicle (sterile 1X PBS) for 18 hours. For *(A,B, H, I, J*) representative pancreatic sections are shown from adult (2 months old), wildtype C57BL/6J mice, n=5 animals. Panel *(C)* shows data from n=11 P5 mice and n=8 p21 mice. Panel *(D)* shows representative images from n=5 wildtype C57BL/6J mouse embryos at indicated stages. For panel *(F)*, data from n=1 fetal, n=2 neonatal and n=3 adult human pancreatic samples is shown. Panel *(G)* shows representative data from (n=5) 2 months old *Ngn3*-Cre:*Rosa26* mTmG mice. *(K) shows* mean of islets from n=5 mice per group split into two pools for each treatment. The error bars represent SEM. \*\*\**P*<0.005, determined by using a two-tailed Student’s *t*-test for (A) and and 1-way ANOVA followed by a Šídák post-hoc test for (K). Scale bar: 50 μm.

**Supplemental Fig. S2.**
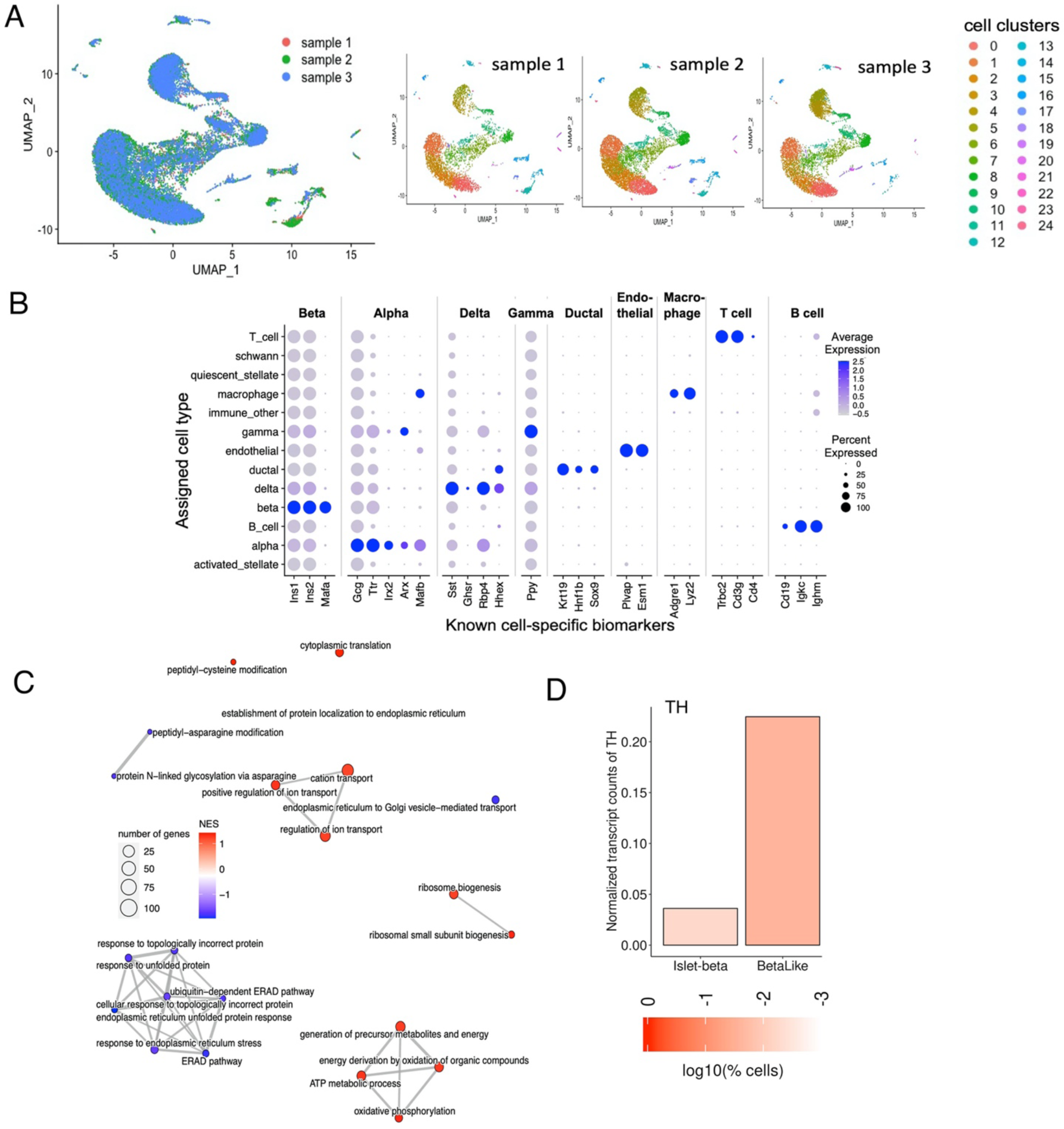
(Related to **Fig. 2).** *(A)* Overlay of the three replicate samples in the integrated dataset and depiction of the three samples individually in the UMAP space. The colors represent the cell clusters as identified by Seurat. *(C)* Enriched pathways clustered into pathway modules based on their number of shared genes. Pathways that share many common genes are clustered together, where the edge thickness is proportional to the Jaccard similarity (fraction of shared genes to the total number of genes in both pathways). Upregulated pathways (positive Normalized Enrichment Score - NES) are colored red, while downregulated pathways (NES, 0) are colored blue. comparison between *Th*+ and *Th*- beta cells of the major beta cell clusters. The enrichment was calculated for Gene Ontology-Biological process (GO-BP) terms. *(D)* Bar graph output of Beta Cell Hub query (Fang et al. 2019; Weng et al. 2020) showing normalized transcripts counts for *TH* in islet-derived or human stem cell-derived beta like cells (islet-beta and beta-like, respectively). For *(D)* data shows representative images of pancreatic sections from adult (2 months old), wildtype C57BL/6J mice are shown, n=5 mice. Scale bar: 50 μm.

**Supplemental Fig. S3.**
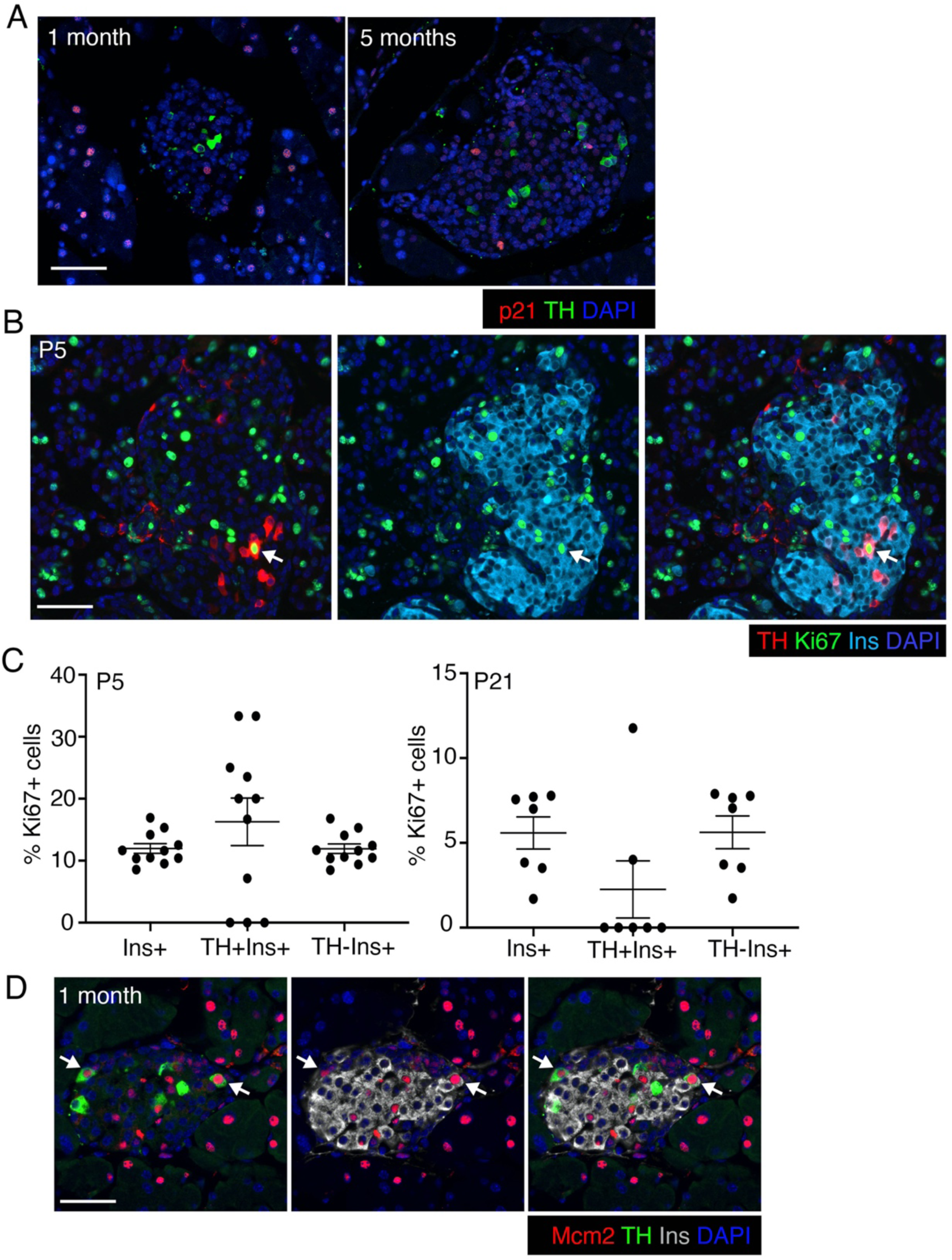
(related to **Fig. 3).** *(A)* Immunofluorescence for senescence marker p21 (red) along with TH (green) in pancreas tissue from adult wildtype mice at indicated ages. *(B)* Immunostaining for replication marker Ki67 (green), along with TH (red) and insulin (Ins: cyan) in P5 pancreatic samples showing the variability of replication of TH-positive beta cells. Nuclei are marked by DAPI in blue. *(C)* Quantification of Ki67 in total, TH+ or TH- beta-cells, expressed as a percentage of total cells in that population, in wildtype P5 and P21 mice. *(D)* Immunostaining for replication marker Mcm2 (red), along with TH green) and insulin (Ins: grey) in representative pancreatic section from 1 month old wildtype C57BL/6J mice shows that TH-positive beta cells can replicate. DAPI labels the nuclei in blue. All panels show representative or mean data from n=5 wildtype C57BL/6J mice. with error-bars showing SEM. \*\*\**P*<0.005, using 1-way ANOVA with Bonferroni post-hoc test. Scale bar: 50 μm.

**Supplemental Fig. S4.**
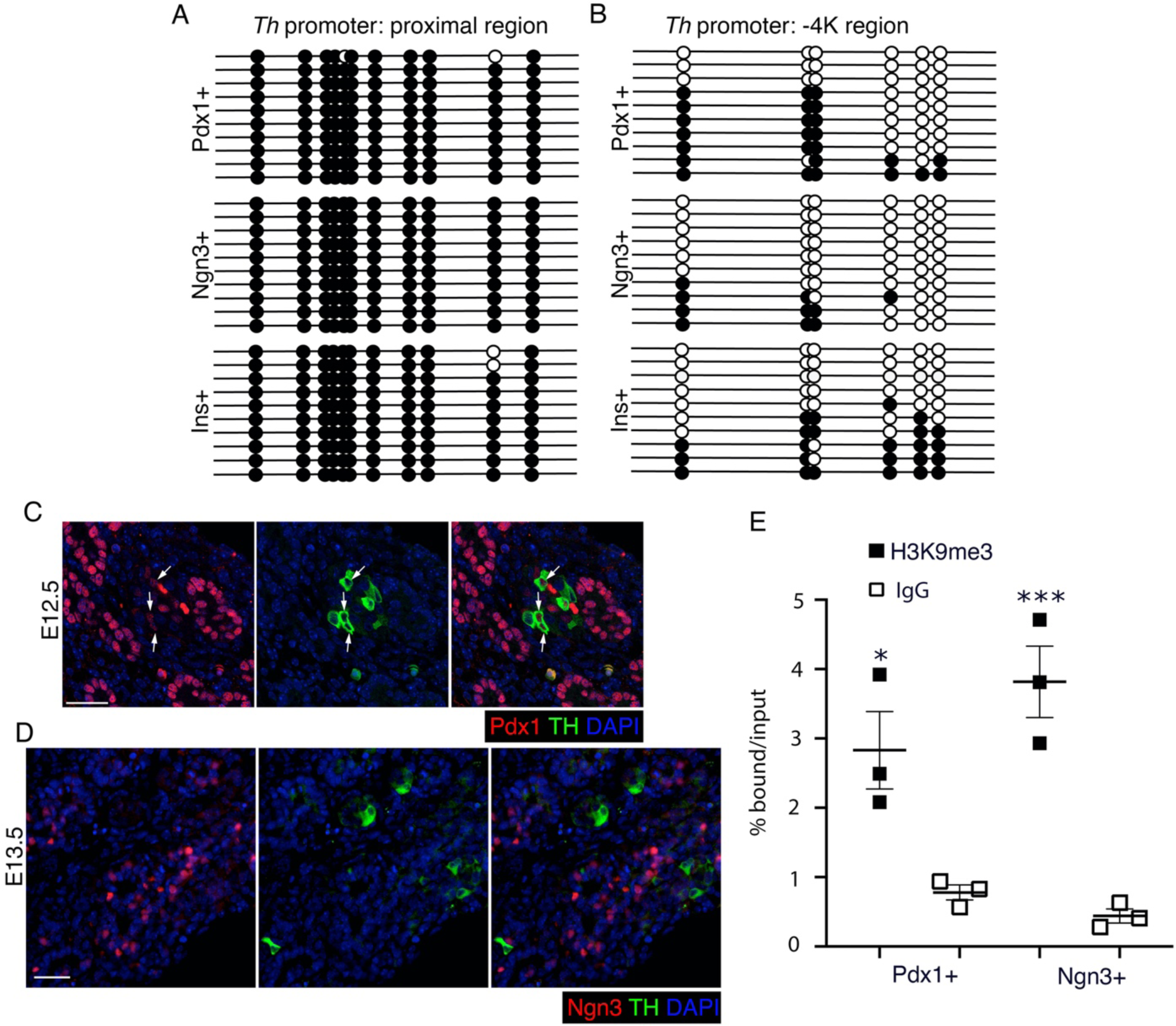
(related to **Fig. 4).** TH promoter undergoes Dnmt3a dependent methylation during the differentiation of endocrine progenitors to beta cells. *(A, B)* Bisulfite sequencing analysis of the proximal *Th* promoter (-21 to -244 bp region) or -5K region of the *Th* promoter, in purified mouse embryonic pancreatic progenitors, endocrine progenitors, and beta cells, showing the methylation of *Th* promoter during beta cell specification. Each horizontal line with dots represents an independent clone and 10 clones are shown here; with filled circles representing a methylated CpG, while an open circle denotes an unmethylated CpG residue. *(C, D)* TH expression (green) analysis using immunofluorescence in the developing wildtype embryonic pancreas; in pancreatic progenitors marked by Pdx1 (red) at E12.5 *(C)*, and endocrine progenitors marked by Ngn3 (red) at E13.5 *(D)*. DAPI marks nuclei in blue. (E) Chromatin immunoprecipitation (ChIP) analysis showing the enrichment for repressive histone modification H3K9me3 (filled squares) or IgG control (open squares) to the -2K region of *Th* promoter in sorted embryonic pancreatic- and endocrine-progenitors. Panels *(A, B)* show representative data from one of the n=3 independent cell preparations, each preparation being a pool of cells derived from multiple embryos. Panels (*C, D)* show representative examples of data from n=5 independent pancreatic samples. Panel *E* shows data as mean of n=3 independent samples per group, each sample being a pool of cells derived from multiple embryos, with error-bars showing SEM of the mean. \*\*\**P*<0.005, using 1-way ANOVA followed by a Bonferroni post-hoc test. Scale bar: 50 μm.

**Supplemental Fig. S5.**
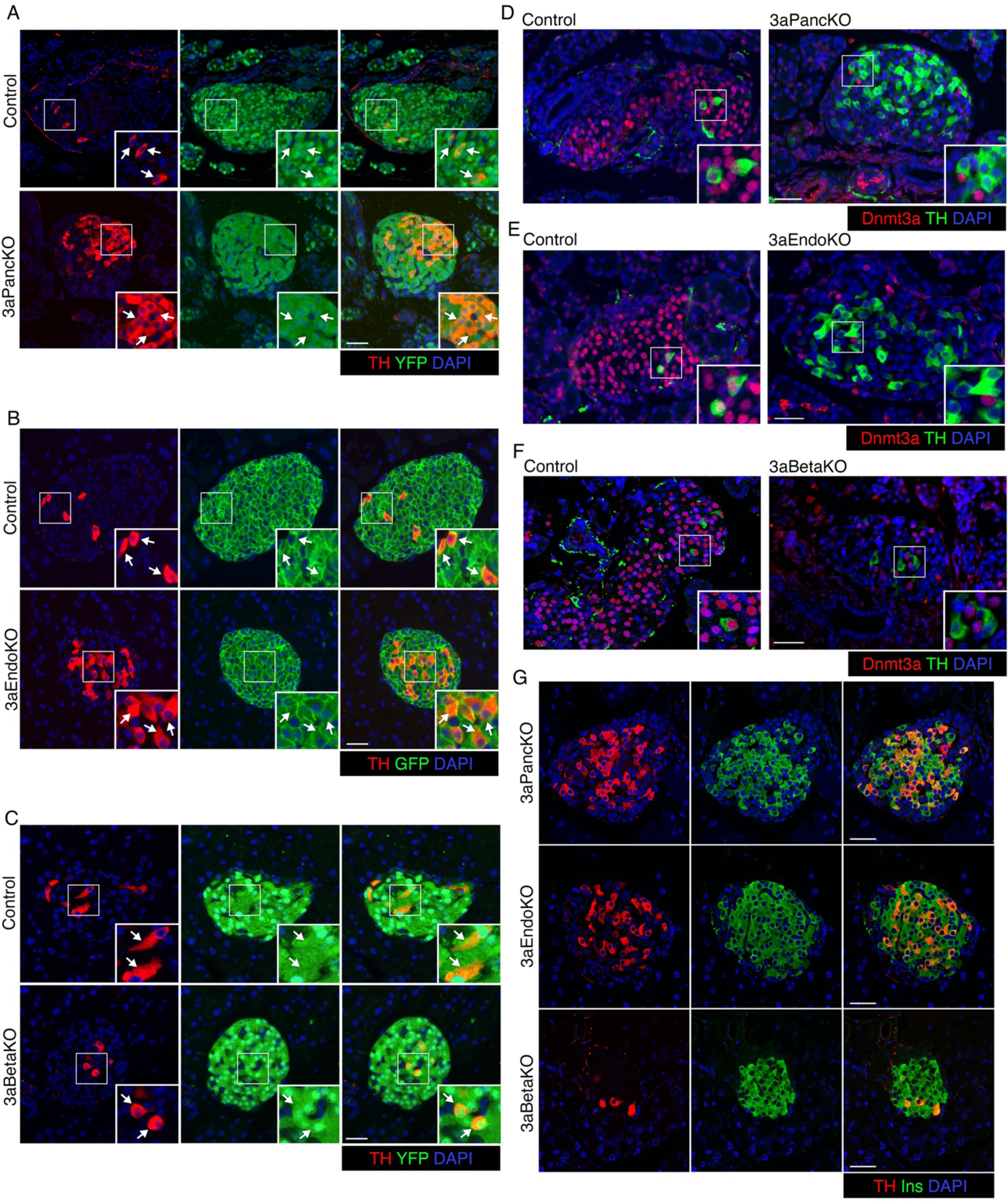
(related to **Fig. 5).** *(A-C)* Immunofluorescence analysis of TH expression (red) along with lineage specific fluorescent label (GFP or YFP: green) in pancreatic sections from adult (2.5 months old) mice with Dnmt3a ablation at various stages of pancreas development and littermate controls, to show efficiency of Cre-recombination as marked by GFP or YFP. Cre recombination in pancreatic progenitor (3aPancKO, *A*) and beta cell lineage (3aBetaKO, *C*) is marked by Cre-driven Rosa26R-*YFP*, while Cre recombination in the endocrine progenitor lineage (3aEndoKO, *B*) is marked by Cre-driven Rosa26R-mTmG. Insets shows a 2X magnified view of examples areas marked by white boxes. *(D-F)* Immunofluorescence analysis of Dnmt3a expression (red) along with TH (green) in pancreatic sections from neonatal (P5) mice to confirm Dnmt3a ablation in 3aPancKO, 3aEndoKO, and 3aBetaKO models. Insets shows a 2X magnified view of examples areas marked by white boxes. We chose neonatal samples to confirm Dnmt3a ablation as Dnmt3a expression declines dramatically after weaning. *(G)* Confocal images of immunolabeling for TH expression (red) along with insulin (Ins: green) in pancreatic sections from adult (2.5 months old) mice with Dnmt3a KO in pancreatic, endocrine, and beta cell lineages, showing co-localization of TH and Ins. All panels show representative immunofluorescence examples from at least n=5 independent pancreatic samples. Scale bar: 50 μm.

**Supplemental Fig. S6.**
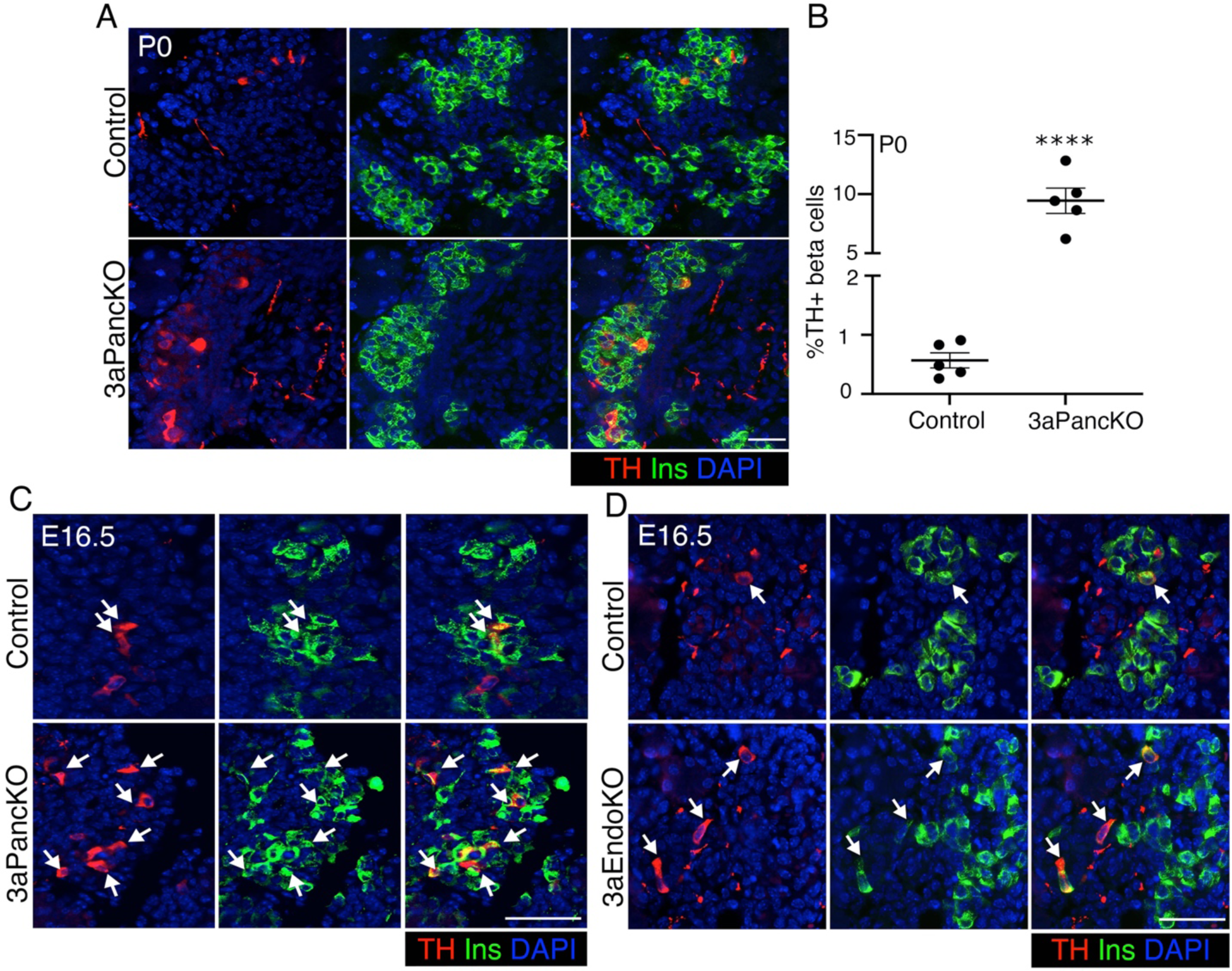
(related to **Fig. 6).** *(A-B)* Immunofluorescence analysis *(A)* and quantification *(B)* of TH expression in pancreatic sections from neonatal mice at P0 with Dnmt3a ablation in pancreatic progenitors (3aPancKO) along with age-matched littermate controls at indicated ages. TH is shown in red, insulin (Ins) in green, and nuclei are labeled by DAPI in blue. Immunofluorescence data shows representative examples from n=5 samples for P0 mice or n=8 samples for P5 mice. *(C, D)* Immunofluorescence analysis of TH expression (red) with insulin (Ins: green) in pancreatic sections from mouse embryos at E16.5 with Dnmt3a ablation in pancreatic (3aPancKO, panel *C*) and endocrine (3aEndoKO, panel *D*) progenitors along with age-matched littermate controls. Nuclei are labeled by DAPI in blue. Arrows mark cells co-expressing TH and insulin. Data shows representative immunofluorescent images and quantification from n=5 KO and littermate control embryos or mice for each group. Error-bars show SEMof the mean. ns= *P* value not significant (>0.05), determined by using a two-tailed Student’s *t*-test. Scale bar: 50 μm.

**Supplemental Fig. S7.**
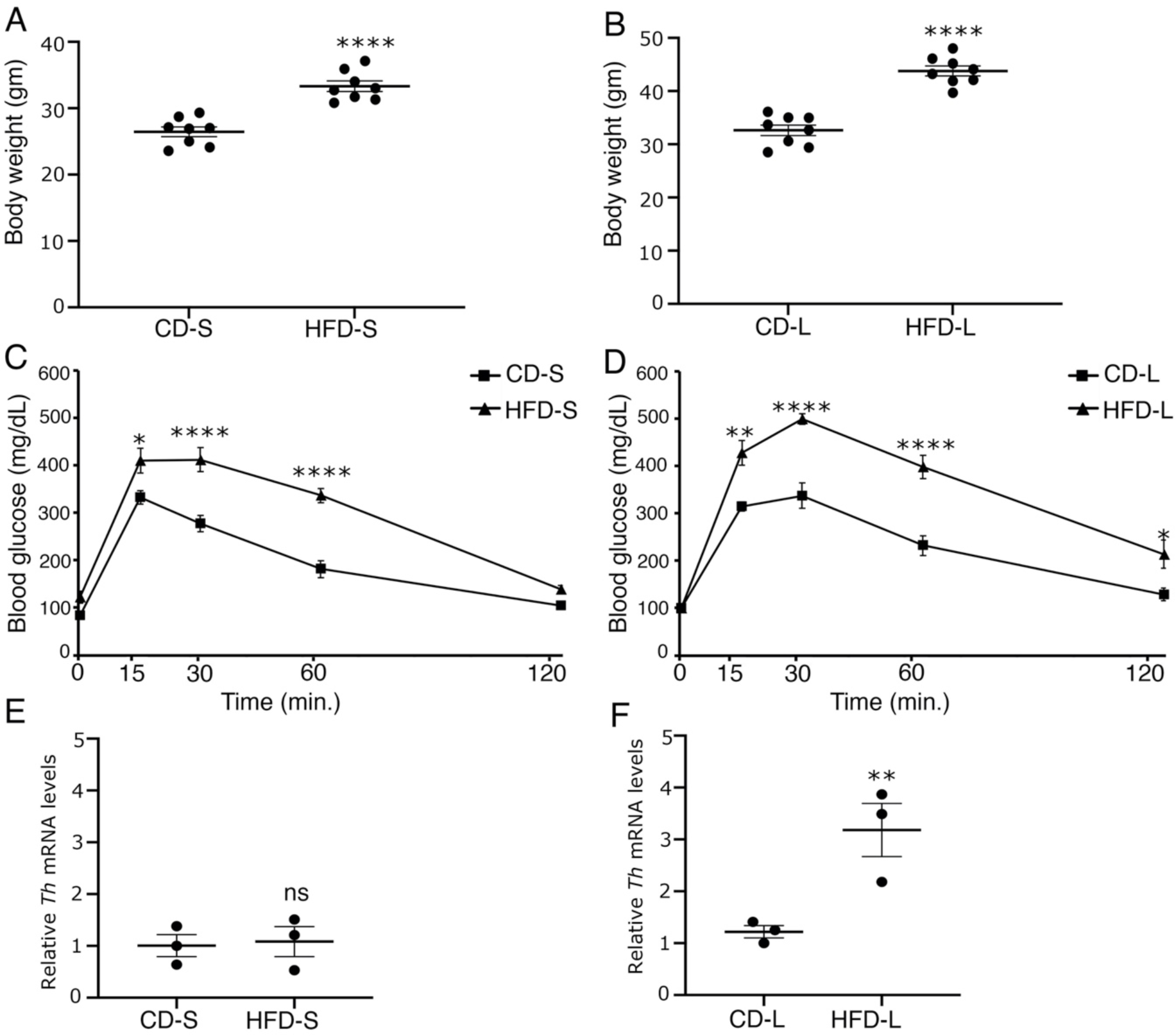
(related to **Fig. 7)**. (*A-D*) Body weight *(A, B)* and intra-peritoneal glucose tolerance tests (IP-GTT) *(C, D)* in adult wildtype 7 weeks old mice fed with a high fat- or control-diet for a short term of 8 weeks (HFD-S or CD-S; panels *A, C*), or long-term for 16 weeks (HFD-L and CD-L; panels *B, D*). *(E, F)* levels of *Th* mRNA relative to housekeeping gene *Cyclophilin*, comparing CD-S vs HFD-S *(E)*, or CD-L vs HFD-L *(F)* cohorts. For *(A-D)*, n=8 animals. For *(E, F),* n=3 animals. The error bars represent SEM. ns= *P* value not significant (>0.05), \**P*<0.05, \*\**P*<0.01, \*\*\**P*<0.005, \*\*\*\**P*<0.001, determined by using a two-tailed Student’s *t*-test for body weight measurements (*A, B)* and mRNA levels *(E, F)*, while 1-way ANOVA followed by Bonferroni post-hoc test was used for IP-GTT *(C, D)*.

## Notes

### Competing Interest Statement

The authors have declared no competing interest.

