## Supplemental Figures and Legends for "DNA methylation Dependent Restriction of Tyrosine Hydroxylase Contributes to Pancreatic *β*-cell Heterogeneity"

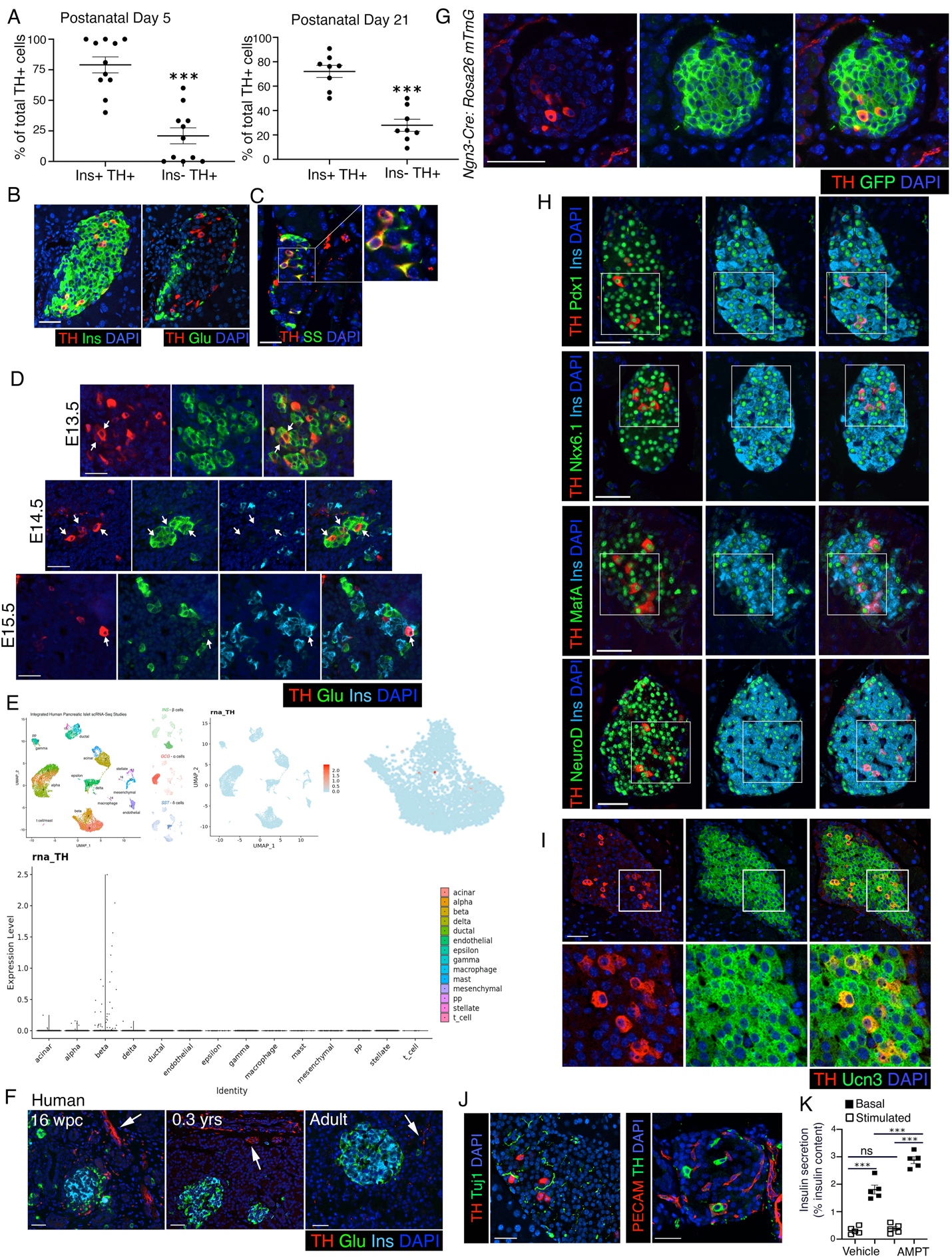

**Supplemental Fig. S1.** *(A)* Immunofluorescence analysis of adjacent pancreatic sections from adult wildtype mice (2 months old) for TH (red) and insulin (Ins: green) in the left panel, and TH (red) and glucagon (Glu: green) in the right panel, showing that TH overlaps with insulin but not glucagon in the adult islets. DAPI counter-stains the nuclei (blue). *(B)* Immunostaining for TH (red) and somatostatin (SS: green), with DAPI (blue), showing that TH also marks a fraction of adult delta cells. Right panel shows a 2X magnified view of the area marked by a white square. *(C)* Morphometric quantification of TH-expressing cells assessed for insulin expression (shown as percentage of total TH-positive cells) at postnatal days 5 (left panel, n=11 wildtype mice) and 21 (right panel, n=8 wildtype mice), showing that majority of TH-expressing cells in the postnatal pancreas are beta cells. *(D)* Immunostaining for Tyrosine Hydroxylase (TH: red), glucagon (Glu: green), and Insulin (Ins: cyan), with DAPI in blue in embryonic pancreas at E13.5, E14.5, and E15.5. Arrows indicate cells co-expressing TH and Glu (top and middle panels; E13.5 and E14.5) or TH and Ins (bottom panel; E15.5). *(E)* Data shows distribution of *TH* mRNA expression in different endocrine cell populations in single-cell RNA sequencing (scRNA-seq) data meta-analyzed and made publicly accessible by (Mawla and Huising 2019) <https://www.huisinglab.com/diabetes_2019/index.html>. The data shown is an integration of four human pancreatic islet datasets, as noted in the study. Top panel shows a UMAP (Uniform Manifold Approximation and Projection) plot, while bottom panel shows a violin plot. *(F)* Immunostaining for TH (red), glucagon (Glu; green), and insulin (Ins; cyan), with DAPI (blue) in fetal (16 weeks p.c./gestation), neonatal (0.3 years) and adult human pancreatic sections, showing absence of TH in human islets. Arrows mark sympathetic nerve fibers that express TH, serving as an internal positive control. *(G)* Representative pancreatic sections from *Ngn3*-Cre:*Rosa26* mTmG lineage reporter mice at 2 months of age, stained for TH (red) and GFP (green) with DAPI in blue. *(H)* Full, un-cropped images corresponding to Fig. 1*(G)* for immunofluorescence analyses showing overlap of TH (red) with key transcription factor hallmarks of beta cell identity, namely, Pdx1 (top left), Nkx6.1 (top right), MafA (bottom left) and NeuroD1 (bottom right) shown in green. Nuclei are marked in blue with DAPI. The regions marked by white box are zoomed 2X and presented in Fig. 1*(G)*. *(I)* Overlap of TH (red) with mature beta cell marker Urocortin3 (Ucn3: green) in adult (2 months old) wildtype pancreas. DAPI (blue) counterstains the nuclei. Inset shows a 2.5X zoomed view of the area marked by white boxes. *(J)* Immunostaining for TH (red) with sympathetic neuronal marker (shown in green) Tuj1 (left) and Nestin (right), showing interaction of sympathetic afferents with TH-positive islet cells. *(K)* Static incubation GSIS in islets from 8 weeks old C57BL/6J mice, pretreated either with 10 μM AMPT or vehicle (sterile 1X PBS) for 18 hours. For *(A,B, H, I, J*) representative pancreatic sections are shown from adult (2 months old), wildtype C57BL/6J mice, n=5 animals. Panel *(C)* shows data from n=11 P5 mice and n=8 p21 mice. Panel *(D)* shows representative images from n=5 wildtype C57BL/6J mouse embryos at indicated stages. For panel *(F)*, data from n=1 fetal, n=2 neonatal and n=3 adult human pancreatic samples is shown. Panel *(G)* shows representative data from (n=5) 2 months old *Ngn3*-Cre:*Rosa26* mTmG mice. *(K) shows* mean of islets from n=5 mice per group split into two pools for each treatment. The error bars represent standard error (SEM) of the mean. ****P*<0.005, determined by using a two-tailed Student’s *t*-test for (A) and and 1-way ANOVA followed by a Šídák post-hoc test for (K). Scale bar: 50 μm.

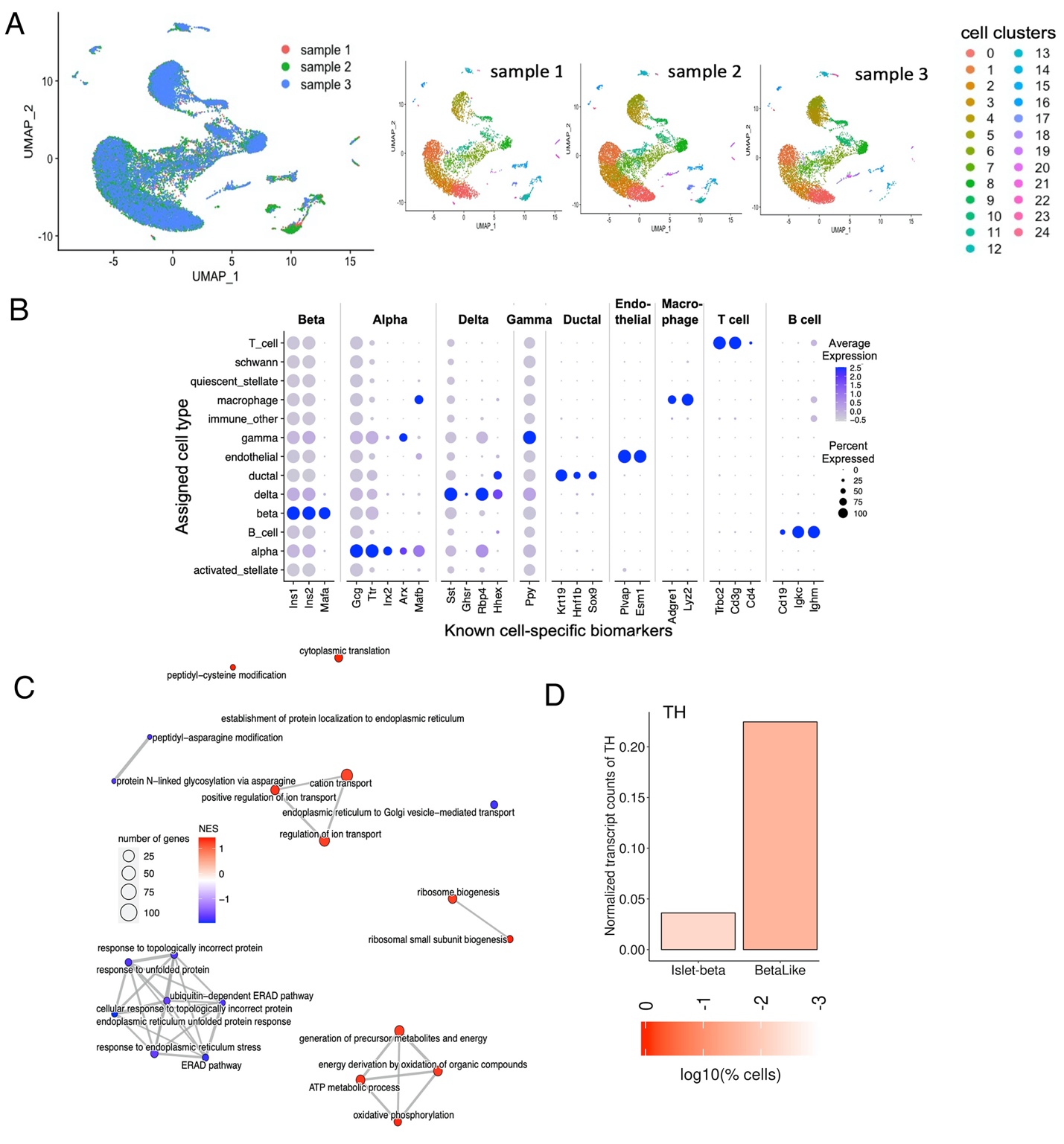

**Supplemental Fig. S2.** *(A)* Overlay of the three replicate samples in the integrated dataset and depiction of the three samples individually in the UMAP space. The colors represent the cell clusters as identified by Seurat. *(C)* Enriched pathways clustered into pathway modules based on their number of shared genes. Pathways that share many common genes are clustered together, where the edge thickness is proportional to the Jaccard similarity (fraction of shared genes to the total number of genes in both pathways). Upregulated pathways (positive Normalized Enrichment Score - NES) are colored red, while downregulated pathways (NES , 0) are colored blue. comparison between *Th*+ and *Th*- beta cells of the major beta cell clusters. The enrichment was calculated for Gene Ontology-Biological process (GO-BP) terms. *(D)* Bar graph output of Beta Cell Hub query (Fang et al. 2019; Weng et al. 2020) showing normalized transcripts counts for *TH* in islet-derived or human stem cell-derived beta like cells (islet-beta and beta-like, respectively). For *(D)* data shows representative images of pancreatic sections from adult (2 months old), wildtype C57BL/6J mice are shown, n=5 mice. Scale bar: 50 μm.

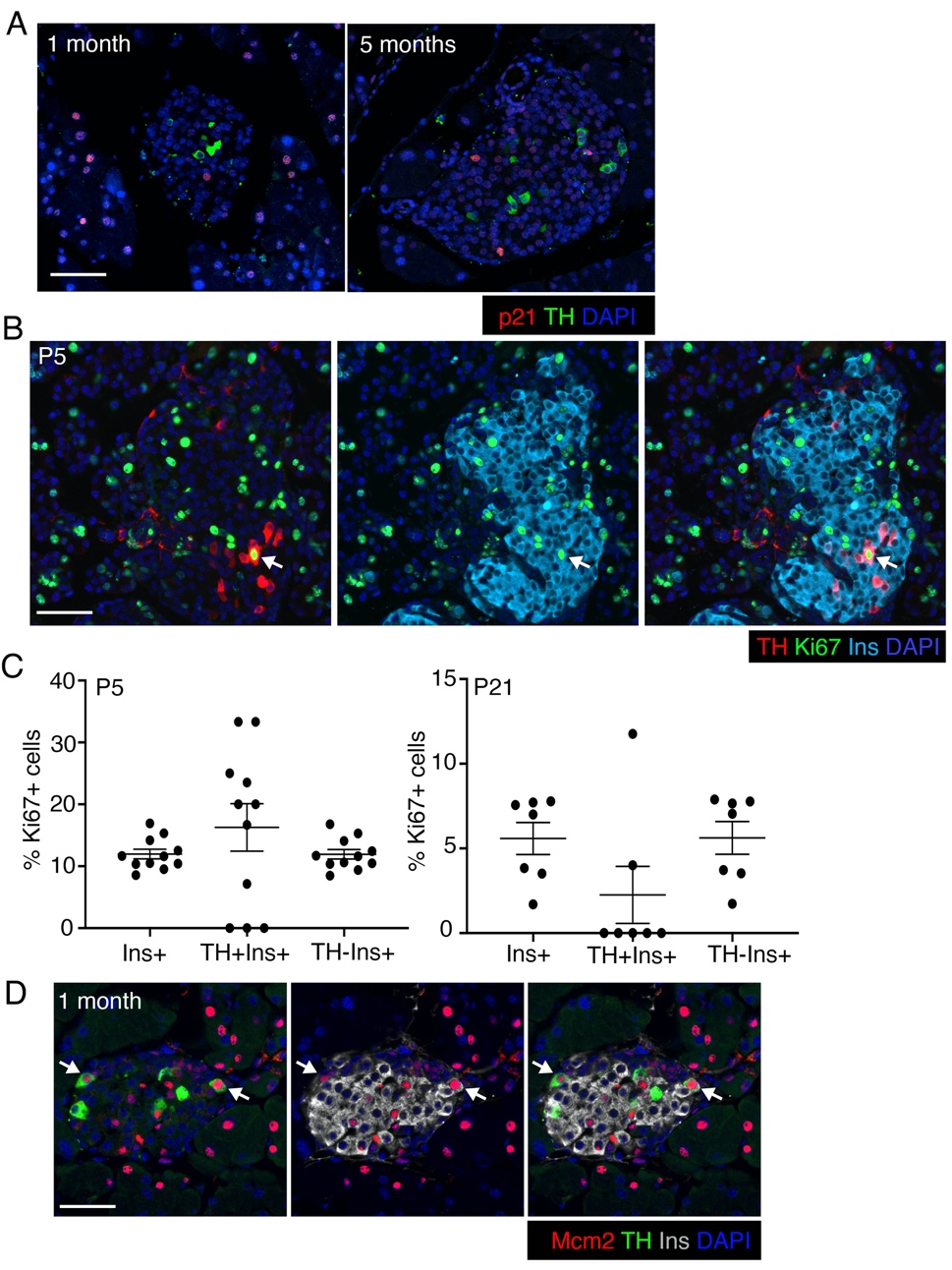

**Supplemental Fig. S3.** *(A)* Immunofluorescence for senescence marker p21 (red) along with TH (green) in pancreas tissue from adult wildtype mice at indicated ages. *(B)* Immunostaining for replication marker Ki67 (green), along with TH (red) and insulin (Ins: cyan) in P5 pancreatic samples showing the variability of replication of TH-positive beta cells. Nuclei are marked by DAPI in blue. *(C)* Quantification of Ki67 in total, TH+ or TH- beta-cells, expressed as a percentage of total cells in that population, in wildtype P5 and P21 mice. *(D)* Immunostaining for replication marker Mcm2 (red), along with TH green) and insulin (Ins: grey) in representative pancreatic section from 1 month old wildtype C57BL/6J mice shows that TH-positive beta cells can replicate. DAPI labels the nuclei in blue. All panels show representative or mean data from n=5 wildtype C57BL/6J mice. with error-bars showing standard error (SEM) of the mean. ****P*<0.005, using 1-way ANOVA with Bonferroni post-hoc test. Scale bar: 50 μm.

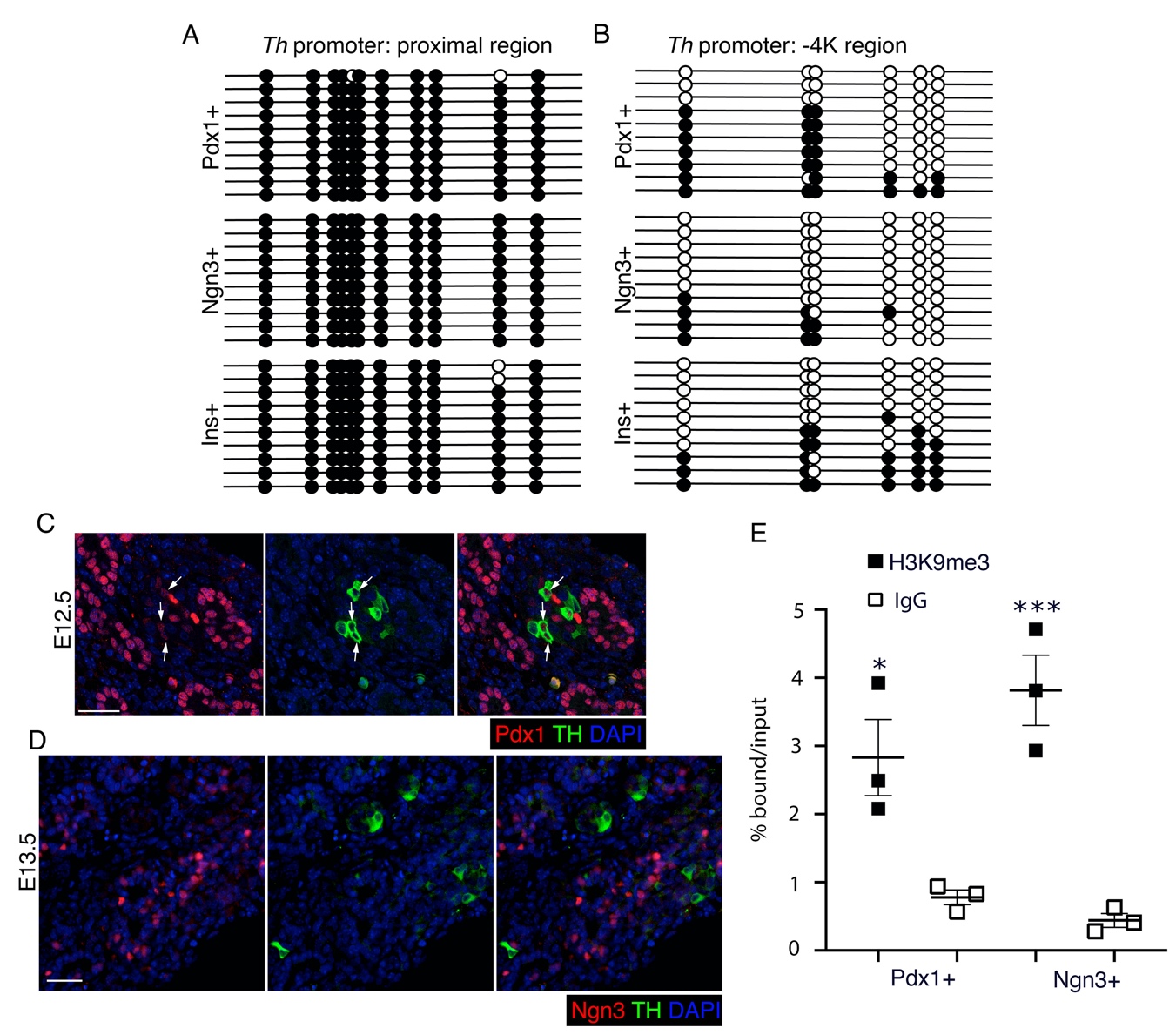

**Supplemental Fig. S4.** TH promoter undergoes Dnmt3a dependent methylation during the differentiation of endocrine progenitors to beta cells. *(A, B)* Bisulfite sequencing analysis of the proximal *Th* promoter (-21 to -244 bp region) or -5K region of the *Th* promoter, in purified mouse embryonic pancreatic progenitors, endocrine progenitors, and beta cells, showing the methylation of *Th* promoter during beta cell specification. Each horizontal line with dots represents an independent clone and 10 clones are shown here; with filled circles representing a methylated CpG, while an open circle denotes an unmethylated CpG residue. *(C, D)* TH expression (green) analysis using immunofluorescence in the developing wildtype embryonic pancreas; in pancreatic progenitors marked by Pdx1 (red) at E12.5 *(C)*, and endocrine progenitors marked by Ngn3 (red) at E13.5 *(D)*. DAPI marks nuclei in blue. (E) Chromatin immunoprecipitation (ChIP) analysis showing the enrichment for repressive histone modification H3K9me3 (filled squares) or IgG control (open squares) to the -2K region of *Th* promoter in sorted embryonic pancreatic- and endocrine-progenitors. Panels *(A, B)* show representative data from one of the n=3 independent cell preparations, each preparation being a pool of cells derived from multiple embryos. Panels (*C, D)* show representative examples of data from n=5 independent pancreatic samples. Panel *E* shows data as mean of n=3 independent samples per group, each sample being a pool of cells derived from multiple embryos, with error-bars showing standard error (SEM) of the mean. ****P*<0.005, using 1-way ANOVA followed by a Bonferroni post-hoc test. Scale bar: 50 μm.

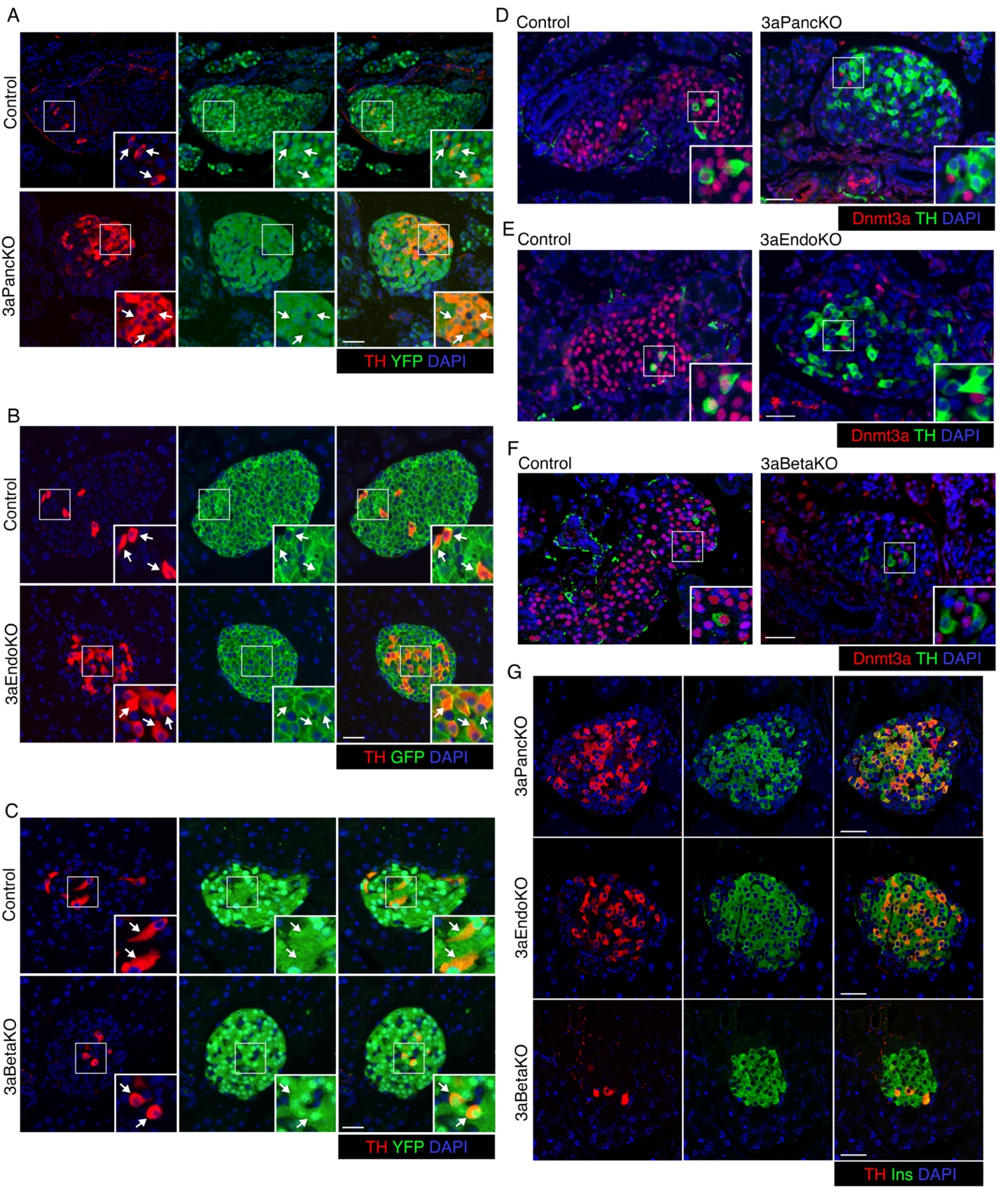

**Supplemental Fig. S5.** *(A-C)* Immunofluorescence analysis of TH expression (red) along with lineage specific fluorescent label (GFP or YFP: green) in pancreatic sections from adult (2.5 months old) mice with Dnmt3a ablation at various stages of pancreas development and littermate controls, to show efficiency of Cre-recombination as marked by GFP or YFP. Cre recombination in pancreatic progenitor (3aPancKO, *A*) and beta cell lineage (3aBetaKO, *C*) is marked by Cre-driven Rosa26R-*YFP*, while Cre recombination in the endocrine progenitor lineage (3aEndoKO, *B*) is marked by Cre-driven Rosa26R-mTmG. Insets shows a 2X magnified view of examples areas marked by white boxes. *(D-F)* Immunofluorescence analysis of Dnmt3a expression (red) along with TH (green) in pancreatic sections from neonatal (P5) mice to confirm Dnmt3a ablation in 3aPancKO, 3aEndoKO, and 3aBetaKO models. Insets shows a 2X magnified view of examples areas marked by white boxes. We chose neonatal samples to confirm Dnmt3a ablation as Dnmt3a expression declines dramatically after weaning. *(G)* Confocal images of immunolabeling for TH expression (red) along with insulin (Ins: green) in pancreatic sections from adult (2.5 months old) mice with Dnmt3a KO in pancreatic, endocrine, and beta cell lineages, showing co-localization of TH and Ins. All panels show representative immunofluorescence examples from at least n=5 independent pancreatic samples. Scale bar: 50 μm.

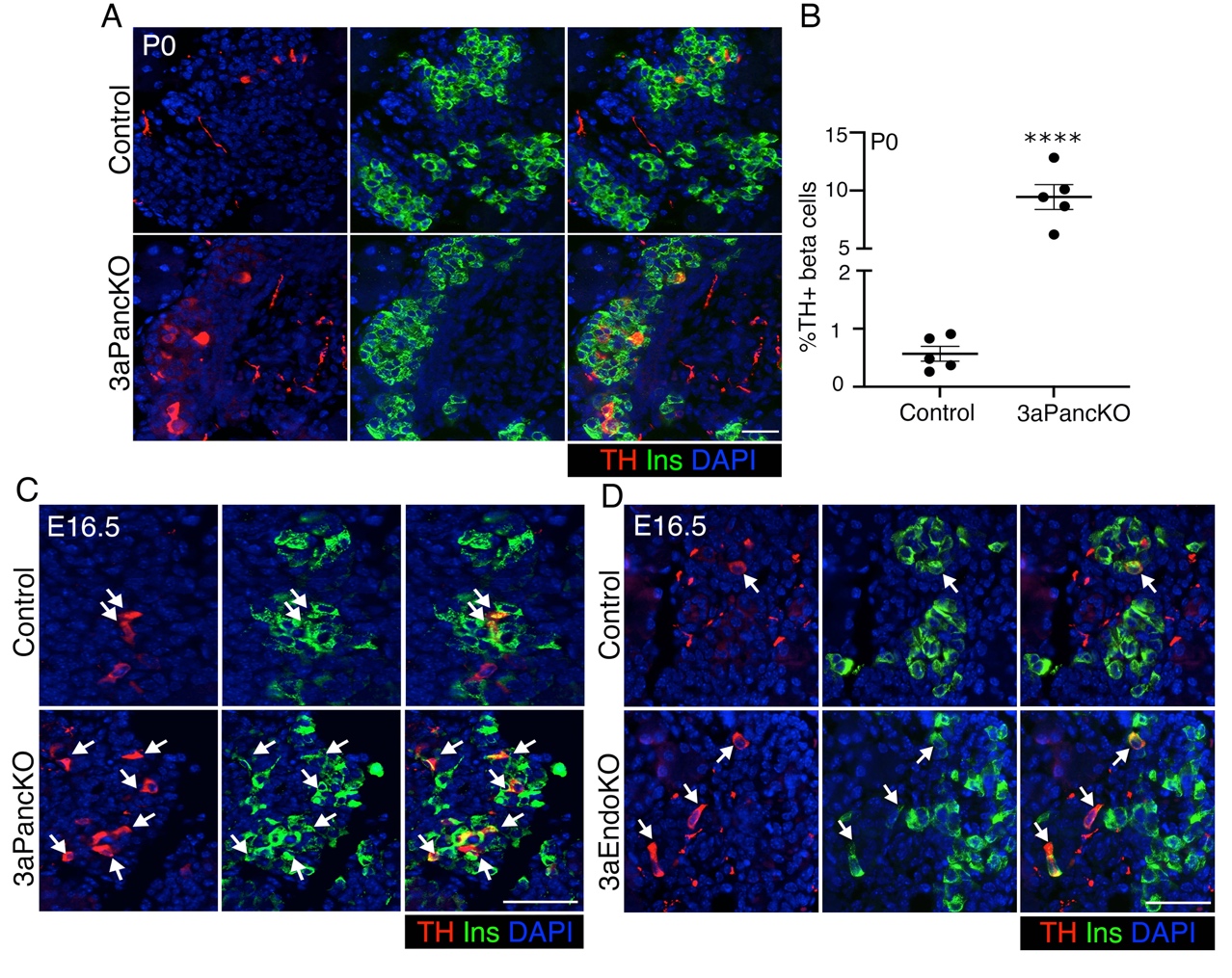

**Supplemental Fig. S6.** *(A-B)* Immunofluorescence analysis *(A)* and quantification *(B)* of TH expression in pancreatic sections from neonatal mice at P0 with Dnmt3a ablation in pancreatic progenitors (3aPancKO) along with age-matched littermate controls at indicated ages. TH is shown in red, insulin (Ins) in green, and nuclei are labeled by DAPI in blue. Immunofluorescence data shows representative examples from n=5 samples for P0 mice or n=8 samples for P5 mice. *(C, D)* Immunofluorescence analysis of TH expression (red) with insulin (Ins: green) in pancreatic sections from mouse embryos at E16.5 with Dnmt3a ablation in pancreatic (3aPancKO, panel *C*) and endocrine (3aEndoKO, panel *D*) progenitors along with age-matched littermate controls. Nuclei are labeled by DAPI in blue. Arrows mark cells co-expressing TH and insulin. Data shows representative immunofluorescent images and quantification from n=5 KO and littermate control embryos or mice for each group. Error-bars show standard error (SEM) of the mean. ns= *P* value not significant (>0.05), determined by using a two-tailed Student’s *t*-test. Scale bar: 50 μm.

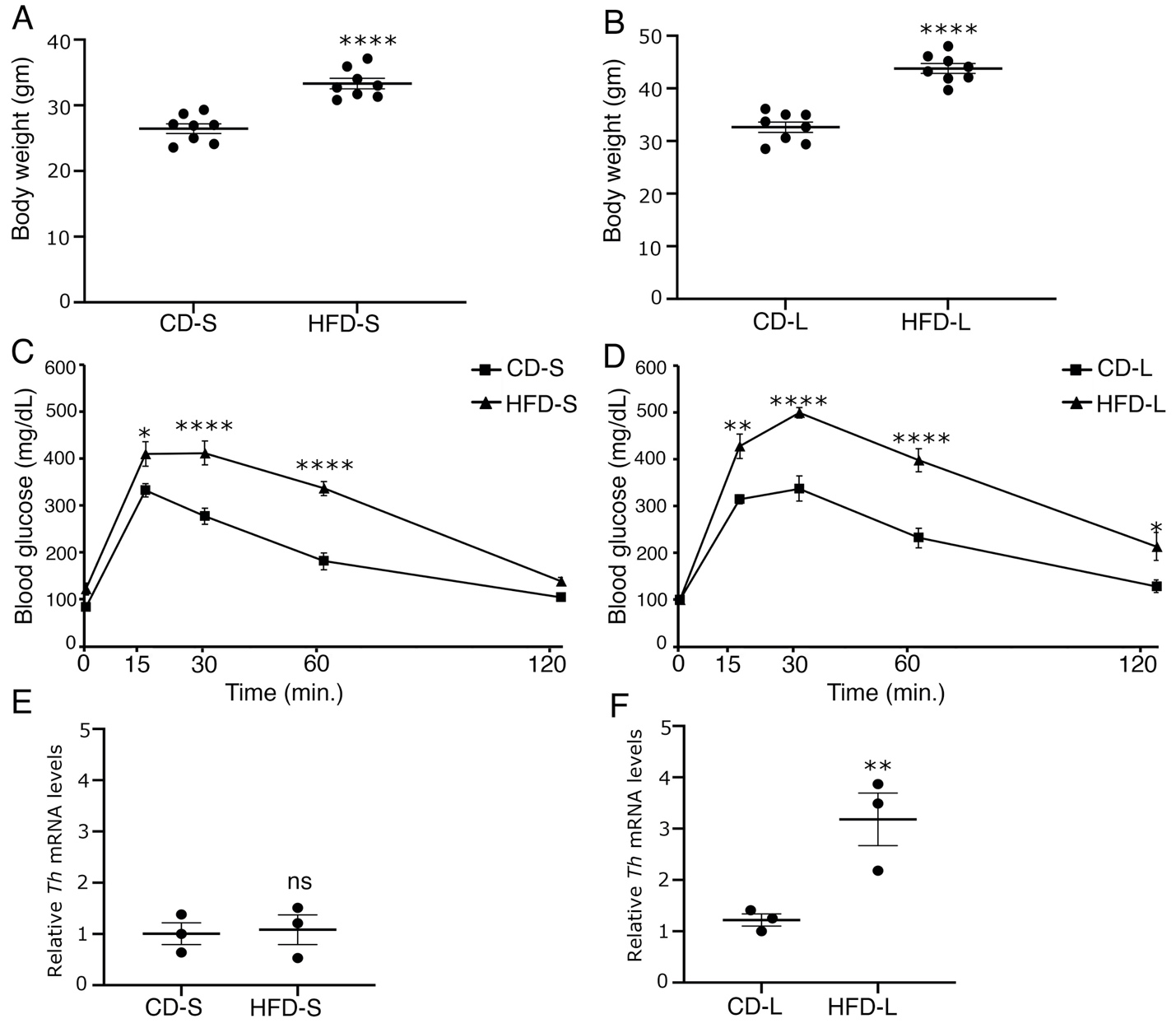

**Supplemental Fig. S7**. (*A-D*) Body weight *(A, B)* and intra-peritoneal glucose tolerance tests (IP-GTT) *(C, D)* in adult wildtype 7 weeks old mice fed with a high fat- or control-diet for a short term of 8 weeks (HFD-S or CD-S; panels *A, C*), or long-term for 16 weeks (HFD-L and CD-L; panels *B, D*). *(E, F)* levels of *Th* mRNA relative to housekeeping gene *Cyclophilin*, comparing CD-S vs HFD-S *(E)*, or CD-L vs HFD-L *(F)* cohorts. For *(A-D)*, n=8 animals. For *(E, F),* n=3 animals. The error bars represent standard error (SEM) of mean. ns= *P* value not significant (>0.05), **P*<0.05, ***P*<0.01, ****P*<0.005, *****P*<0.001, determined by using a two-tailed Student’s *t*-test for body weight measurements (*A, B)* and mRNA levels *(E, F)*, while 1-way ANOVA followed by Bonferroni post-hoc test was used for IP-GTT *(C, D)*.
