## Supplemental Tables for "DNA methylation Dependent Restriction of Tyrosine Hydroxylase Contributes to Pancreatic *β*-cell Heterogeneity"

**Supplemental Table 1**

**Clinical Characteristics of human subjects (Organ donors)**

Fasting Plasma Glucose not available, C-peptide levels are shown where available.

| **Status** | **Age (Year)** | **Sex** | **BMI** | **C-peptide**  **ng/mL** |
| --- | --- | --- | --- | --- |
| Fetus  Non-diabetic | 16 weeks p.c. | - | - | - |
| Fetus  Non-diabetic  nPOD 6201 | 36 weeks | M | - | - |
| Neonatal  Non-diabetic  nPOD 6183 | 0.3 years | M | - | - |
| Adult  Non-diabetic  nPOD 6057 | 22 years | M | 26 | 16.23 |
| Adult  Non-diabetic  nPOD 6003 | 23 years | F | 29.3 | Not available |
| Adult  Non-diabetic  nPOD 6004 | 33 years | M | Not known | Not available |

**Supplemental Table 2**

**Antibodies used for immunofluorescence analyses**

| **Antibody** | **Dilution** | **Source and Catalog Number** |
| --- | --- | --- |
| Mouse anti-Glucagon | 1:1000 | Sigma-Aldrich G2654-.2ML |
| Rabbit anti-Glucagon | 1:500 | Immunostar 20076 |
| Guinea pig anti-Insulin | 1:800 | Abcam 195956 |
| Rabbit anti-insulin | 1:20,000 | Abcam ab181547 |
| Rat anti-somatostatin | 1:300 | Millipore-Sigma MAB354 |
| Mouse anti-Nkx6.1 | 1:50 | Developmental Studies Hybridoma Bank F55A1-0S |
| Mouse anti-Pdx1 | 1:200 | Developmental Studies Hybridoma Bank F109-D12-S |
| Rabbit anti-MafA | 1:500 | Bethyl Laboratories A300-611A |
| Goat anti-NeuroD1 | 1:200 | R&D Systems AF2746 |
| Mouse anti-Ki67 | 1:100 | BD Biosciences 550609 |
| Rabbit anti-pHH3 | 1:200 | Millipore-Sigma 06-570 |
| Mouse anti-Mcm2 | 1:500 | BD Biosciences 610701 |
| Rabbit anti-p21 | 1:500 | Abcam ab188224 |
| Mouse anti-Dnmt3a | 1:200 | Novus Biologicals NB120-13888 |
| Rabbit anti-Ucn3 | 1:1000 | Gift from Dr. Mark O. Huising, UC Davis |
| Rabbit anti-Ero1lb | 1:1500 | Gift from Dr. David Ron, University of Cambridge |
| Mouse anti-TH | 1:200 | Millipore-Sigma MAB318 |
| Rabbit anti-TH | 1:500 | Abcam ab112 |
| Chicken anti-TH | 1:200 | Abcam ab76442 |
| Mouse anti-Tuj1 | 1:500 | Biolegend 801201 |
| Chicken anti-GFP | 1:250 | Aves Labs Inc. 1020 |
| Goat anti-PECAM1 | 1:200 | R&D Systems AF3628 |

**Supplemental table 3**

**Antibodies used for ChIP analyses**

| **Antibody** | **Source and Catalog Number** |
| --- | --- |
| Mouse anti-Dnmt3a | Novus Biologicals NB120-13888 |
| Rabbit anti-H3K9me3 | Millipore-Sigma 07-442 |
| Mouse IgG control | Diagenode KCH-819-015 |
| Rabbit IgG control | Diagenode KCH-504-250 |

**Supplemental Table 4**

**Primers used for ChIP analysis**

| **Region** | **Location** | **Forward** | **Reverse** |
| --- | --- | --- | --- |
| *Th* | -2250 to  -1981 bp. | 5’-CCCACCTAGCTTCTGTTGCA-3’ | 5’-CAGGGTAGGACGGAGCTGTA-3’ |
| *C (Arx) Negative control* | +1348 to +1470 bp. | 5’-AGTGCCCCTCTTGCTACCTT-3’ | 5’-TAGGGTGGGGCAAATTTTTA-3’ |

**Supplemental Table 5**

**Primers used for Bisulfite sequencing analysis**

| **Region** | **Location** | **Forward** | **Reverse** |
| --- | --- | --- | --- |
| *Th proximal promoter* | -215 to +9 bp. | 5’- GAGGGTGATTTAGAGGTAGGTGTT-3’ | 5’- TATCTCCACAACCCTTACCAAAC -3’ |
| *Th -2K* | -2250 to  -1981 bp. | 5’-GAGTTTAGATATAGATAAAGGTTTGGAGAG-3’ | 5’- ATAAAACAAACCCAAACTAAATTCC-3’ |
| *Th -4K* | -4691 to  -4488 bp. | 5’-AGGGTTTAGAGTTAGGGTTGAGATA-3’ | 5’-ATCAAAACCAAATACCAAAACAATTA-3’ |

**Supplemental Table 6**

**Primers used for Real-time qRT-PCR**

| **Gene** | **Forward** | **Reverse** |
| --- | --- | --- |
| *Dnmt1* | 5’- caaatagatccccaagatccag-3’ | 5’- cggaactaggtgaagtttcaaaaa-3’ |
| *Th* | 5’- TGTTGGCTGACCGCACAT-3’ | 5’- GCCCCCAGAGATGCAAGTC-3’ |
| *Cyclophilin* | 5’-GTTGGCCAGGCTGGTGTCCAG-3’ | 5’-CTGTGATGAGCTGCTCAGGGTGG-3’ |
